# A novel broad-spectrum antibiotic targets multiple-drug-resistant bacteria with dual binding targets and no detectable resistance

**DOI:** 10.1101/2025.01.11.632532

**Authors:** Wenyan He, Xueting Huan, Yinchuan Li, Qisen Deng, Tao Chen, Wen Xiao, Yijun Chen, Lingman Ma, Nan Liu, Zhuo Shang, Zongqiang Wang

**Affiliations:** State Key Laboratory of Natural Medicines, China Pharmaceutical University, Nanjing, Jiangsu 211198, China; School of Pharmacy, China Pharmaceutical University, Nanjing, Jiangsu, 211198, China; School of Life Science and Technology, China Pharmaceutical University, Nanjing, Jiangsu, 211198, China; School of Pharmaceutical Sciences, Shandong University, Jinan, Shandong 250012, China

## Abstract

The rapid emergence of difficult-to-treat multidrug-resistant pathogens, combined with the scarcity of antibiotics possessing novel mechanisms, poses a significant threat to global public health. Here, we integrated the synthetic-bioinformatic natural product approach with peptide optimization to unveil the antibiotic-producing potential of *Paenibacillaceae* bacteria. Our culture-independent approach led to the discovery of paenimycin, a novel 11-mer *depsi*-lipopeptide featuring an unprecedented dual-binding mechanism. By sequestering the phosphate and hydroxyl groups of lipid A in Gram-negative bacteria, as well as the phosphate groups of teichoic acids in Gram-positive bacteria, paenimycin exhibited potent and broad-spectrum efficacy against MDR pathogens *in vitro* and *in vivo* models. Remarkably, paenimycin demonstrates no detectable resistance, favorable pharmacokinetics and low nephrotoxicity, positioning it as a promising candidate for treating serve and urgent MDR infections.

## Introduction

Over the past century, antibiotics have revolutionized the treatment of infectious diseases and saved billions of lives (*1*). However, the rapid emergence and spread of multidrug-resistant (MDR) bacteria posed a significant threat to global public health and resulted in approximately 1.14 million deaths worldwide in 2021 (*2, 3*). There is an urgent need to develop novel antibiotic featuring distinct mode of action to treat MDR bacteria. The *Paenibacillaceae* family of bacteria, renowned as the source of pivotal antibiotics such as colistin, cilagicin, tyrocidine, and gramicidin, represents a treasure trove of invaluable antimicrobial natural products. To date, over 39 types of antimicrobial compound have been isolated and characterized from this family using traditional culture-dependent approach (Fig. 1A and fig. S1). Recent advancements in genomic sequencing and bioinformatic analysis have revealed that most biosynthetic gene clusters (BGCs) within *Paenibacillaceae* remain uncharacterized. The silence and extremly low yields of these BGCs under laboratory cultivation conditions (culture-dependent approach) hindered the isolation of new antibiotics from this family (*4*).

**Fig. 1.**
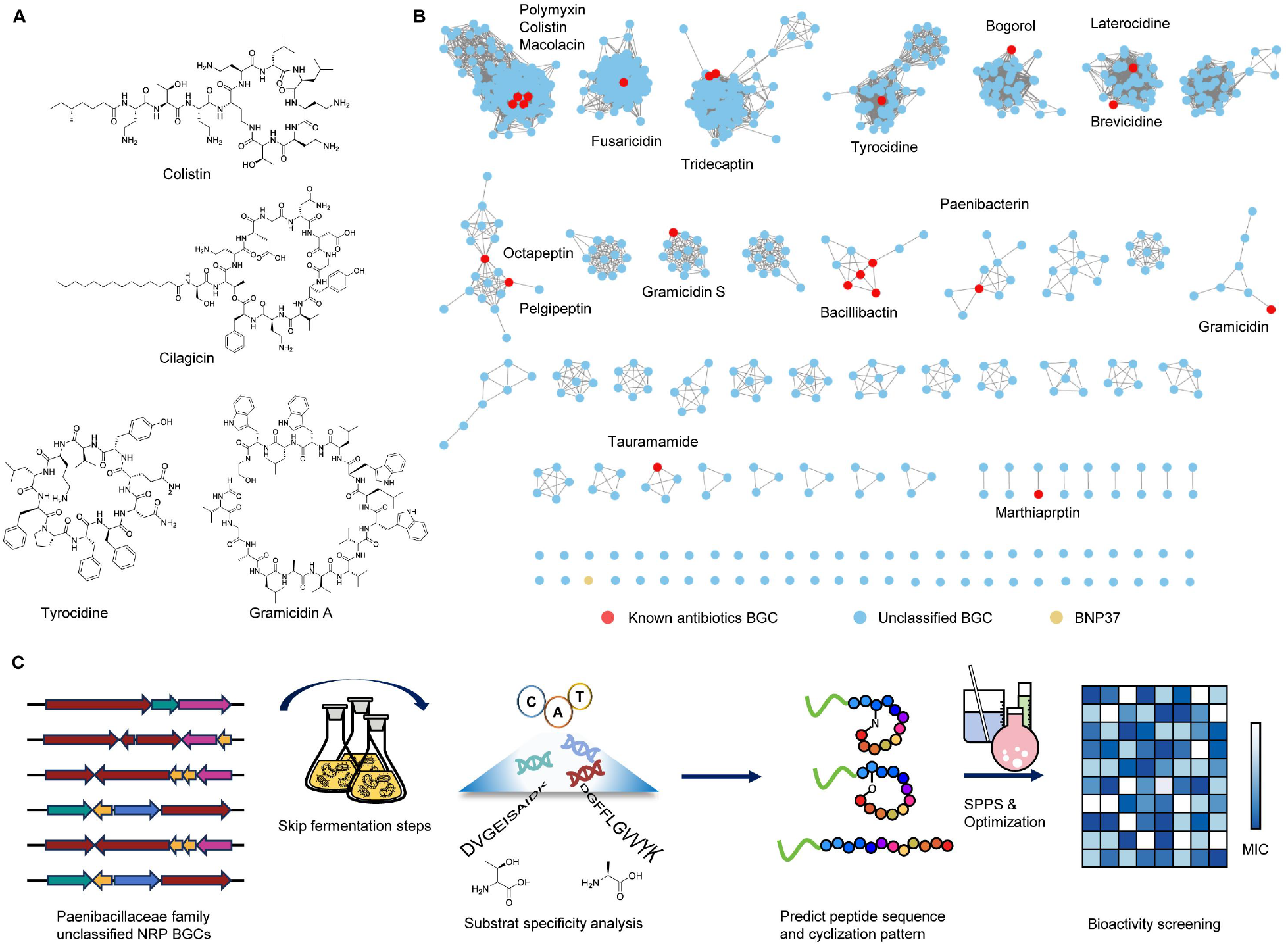
Discovery of *pae* biosynthetic gene cluster from *Paenibacillaceae* family. A) Representative antibiotics identified within *Paenibacillaceae* family. B) Sequence similarity network analysis of all NRP biosynthetic gene clusters (BGCs) of *Paenibacillaceae* family. Known antibiotic BGCs are highlighted in red, unclassified BGCs in blue, and paenimycin BGC is indicated in yellow. C) Overview of the culture-independent antibiotic discovery approach utilized in the present present study. Note: SPPS, Solid phase peptide synthesis.

In recent years, advances in understanding the biosynthetic machinery of microbial-derived secondary metabolites have made it increasingly feasible to predict their structures directly from BGC information with high accuracy (*5, 6*). This allows structure prediction without requiring the expression of BGC or reliance on spectroscopy-based structure elucidation. Currently, the biosynthesis of two major families of peptide secondary metabolites—non-ribosomal peptides (NRPs) and ribosomally synthesized and post-translationally modified peptides (RiPPs)—is particularly well understood (*7, 8*). Their overall structures can be predicted based on adenylation domain specificity for NRPs and precursor peptide sequences for RiPPs and synthesized directly using chemical reaction (i.e., synthetic bioinformatic natural product (synBNP)) (*9*). This approach integrates predictive algorithms with the feasibility of peptide synthesis, bypassing the need for in vivo expression and addressing the challenges posed by silent BGCs (*10ߝ13*) (Fig. 1C).

Herein, we integrated the synBNP approach with peptide structure optimization to systematically investigate the antimicrobial capabilities of NRPs encoded by bacteria in the family of *Paenibacillaceae*. This strategy led to the discovery of a novel broad-spectrum antibiotic, paenimycin, which exhibited potent activity against both Gram-positive bacteria and Gram-negative bacteria *in vitro* and *in vivo*. Mechanistic studies showed that paenimycin uniquely targets two sites: lipid A in Gram-negative bacteria and teichoic acids in Gram-positive bacteria. Unlike colistin, which targets phosphate group of lipid A (*14, 15*), paenimycin binds to a different site on lipid A, enabling it to maintain potent efficacy against colistin-resistant pathogens in both *in vitro* and *in vivo* models. Notably, in contrast to colistin, which is associated with well-documented nephrotoxicity (*16, 17*), paenimycin exhibits significantly reduced *in vivo* nephrotoxicity. Given its potent and broad-spectrum antibacterial activity, unique dual mode of action and low nephrotoxicity, paenimycin emerges as a promising candidate for the development of new antimicrobial therapies to treat complex infections caused by MDR pathogens.

### Discovery of paenimycin

To systematically explore the antimicrobial secondary metabolites encoded by the bacteria in *Paenibacillaceae* family, we retrieved 1245 genomes from public databases along with 11 genomes from our in-house collection. Our study focuses on NRPs, the predominant class of antimicrobial peptides produced by the *Paenibacillaceae* family. AntiSMASH analysis of these genomes identified 17,037 BGCs, which were subsequently deduplicated based on type of BGC, adenylation domain specificity and further refined to include only complete BGCs with 5 to 15 adenylation domain. This process led to the identification of 901 NRP BGCs, which were grouped into 45 clusters and 27 single dots in a sequence similarity network (SSN) analysis using a similarity cutoff of 0.4 (Fig. 1B). We then prioritized 48 representative BGCs from the clusters and singletons, focusing exclusively on those not associated with any known NRP BGCs, thereby ensuring their novelty. To obtain the products of the prioritized NRP BGCs, we employed the synBNP approach involving peptide sequence prediction and solid-phase peptide synthesis. This culture-independent method overcomes the low expression or silence of BGCs typically encountered in culture-dependent antibiotic discovery. The NRP structure prediction was based on the following criteria: (i) the amino acids activated by each adenylation domain in NRPS modules are predicted in silico using 10 signature codes (*18, 19*); (ii) the presence of an epimerization (E) domain indicates the incorporation of D-amino acids (*20*); (iii) the order of NRPS modules corresponds to the amino acid sequence in the peptide (*21*); (iv) the presence of a condensation starter (Cs) domain suggests a lipid chain forms an acyl group with the N -terminus of the peptide core (*22*). However, genomic analysis cannot predict the type of acyl group or the topology of NRPs. Since myristic acid is one of the most common lipids found in bacterial NRPs, all predicted lipopeptides were chemically synthesized with their N-terminal acylated by myristic acid (*12*). NRPs can exist in either linear or cyclized forms. In the cyclized forms, the C-terminal carboxyl group may form an amide bond with either the terminal amino group (head-tail cyclization) or with the side-chain amino group of basic residues (e.g., lysine, 2,4-diaminobutyric acid) (branch-to-tail cyclization). Alternatively, it may form an ester bond with hydroxylated residues (e.g., threonine or serine) in a branch-to-tail topology. To explore all potential structural possibilities, we synthesized the three major topological patterns—linear, head-to-tail cyclization, and branch-to-tail cyclization—for preliminary activity screening. The predicted peptides were chemically synthesized using the gold-standard solid-phase method with HBTU/PyBOP as the coupling agent. This approach yielded a collection of 74 purified peptides predicted from 48 representative NRP BGCs (table S2).

We next screen the antimicrobial activity of 74 synthetic peptide against clinically significant and challenging MDR ESKAPE pathogens (i.e., *Enterococcus faecium, Staphylococcus aureus, Klebsiella penomoniea, Acinetobacter baumannii, Pseudomonas aeruginosa* and *Enterobacter cloacae*), which account for a large proportion of the resistant infections encountered in clinical medicine (fig. S2). Notably, the BNP37 peptide family, comprising five variants (i.e. BNP37L, BNP37C1, BNP37C2, BNP37C3, BNP37C4) with identical sequences but different topologies, exhibited inhibitory activity against ESKAPE pathogens. Their minimum inhibitory concentrations (MICs) ranged from 2 to 64 μg/mL, depending on the specific peptide and targeted strain (Fig. 2B). The structures of the BNP37 peptides were predicted based on the adenylation domain specificity of the *pae* cluster, which is exclusively present in the genome of *Paenibacillus caseinilyticus* GW78 (Fig. 2A). The *pae* cluster comprises three NRPS genes (*pae-B, pae-C, pae-D*), two transporter genes (*pae-E* and *pae-F*), and one regulator gene (*pae-A*). The three NRPS genes together encode an 11 mer peptide predicted to consist of five proteinogenic amino acids (L-Thr, 2× L-Leu, L-Val, L-Asp) and six non-proteinogenic amino acids (3× L-Dab, D-Dab, D-Phe, D-Tyr) with a confidence level of at least 80%. Furthermore, the presence of Cs domain suggests the *pae* cluster encodes lipopeptides. Attempts to isolate the natural peptides encoded by the *pae* cluster from *P. caseinilyticus* GW78 using various media were unsuccessful, likely due to the cluster being silent under these growth conditions. Consequently, we chemically synthesized five BNP37 lipopeptides, one linear form (BNP37L) and four cyclized forms (BNP37C1, BNP37C2, BNP37C3, BNP37C4), and all N-acylated with myristic acid. Among the five variants, the linear peptide BNP37L exhibited the most potent antibacterial activity (MIC 2-4 μg/mL) but showed strong serum binding affinity (Fig. 2B), which may limit its *in vivo* efficacy. In contrast, the cyclized peptide BNP37C2 maintained comparable activity against most ESKAPE strains (MIC 2-8 μg/mL) while showing reduced serum binding ability (Fig. 2B). To enhance druggability, we synthesized 37 BNP37C2 analogues with diverse lipid chains differing in chain length, degree of branching, and substitutions (fig. S3). Antimicrobial and cytotoxicity evaluations revealed that none of these analogues outperformed BNP37C2, which features myristic acid as the acyl substituent (fig. S4A). BNP37C2 demonstrated an optimal balance between antimicrobial activity, serum binding affinity, and cytotoxicity, making it the most promising candidate for further optimization. To assess the contribution of individual amino acid residues to the antibacterial activity of BNP37C2, we synthesized ten peptide variants, each with a single amino acid replaced by alanine. Thr-2, critical for cyclization, was excluded from this analysis (fig. S3 and S4B). Substituting the Dab residues at positions 1, 3, 5, and 8 with alanine resulted in a 2-to 16-fold increase in MIC values. These results highlight the pivotal role of the positively charged Dab residues in maintaining the antibacterial activity of BNP37C2. Considering that BNP37C2 inherently contains four positively charged residues, we investigated the effect of adding extra cationic Dab residues on its activity. Remarkably, substituting the nonpolar D-Tyr residue at position 9 with a D-Dab residue reduced the MIC against carbapenem-resistant *Acinetobacter baumannii* (CRAB) and methicillin resistant *Staphylococcus aureus* (MRSA) by 2-fold and eliminated serum binding (Fig. 2C). This optimized structure, derived from targeted modifications of BNP37C2, has been named paenimycin. Paenimycin is a depsi-lipopeptide consisting of 11 amino acids, including five positively-charged Dab residues and five proteinogenic amino acids (Fig. 2D). The peptide is cyclized via an ester bond between the hydroxyl group of threonine at position 2 and the α-carboxyl group of the terminal aspartic acid, with an N-terminal myristic acid anchoring the structure.

**Fig. 2.**
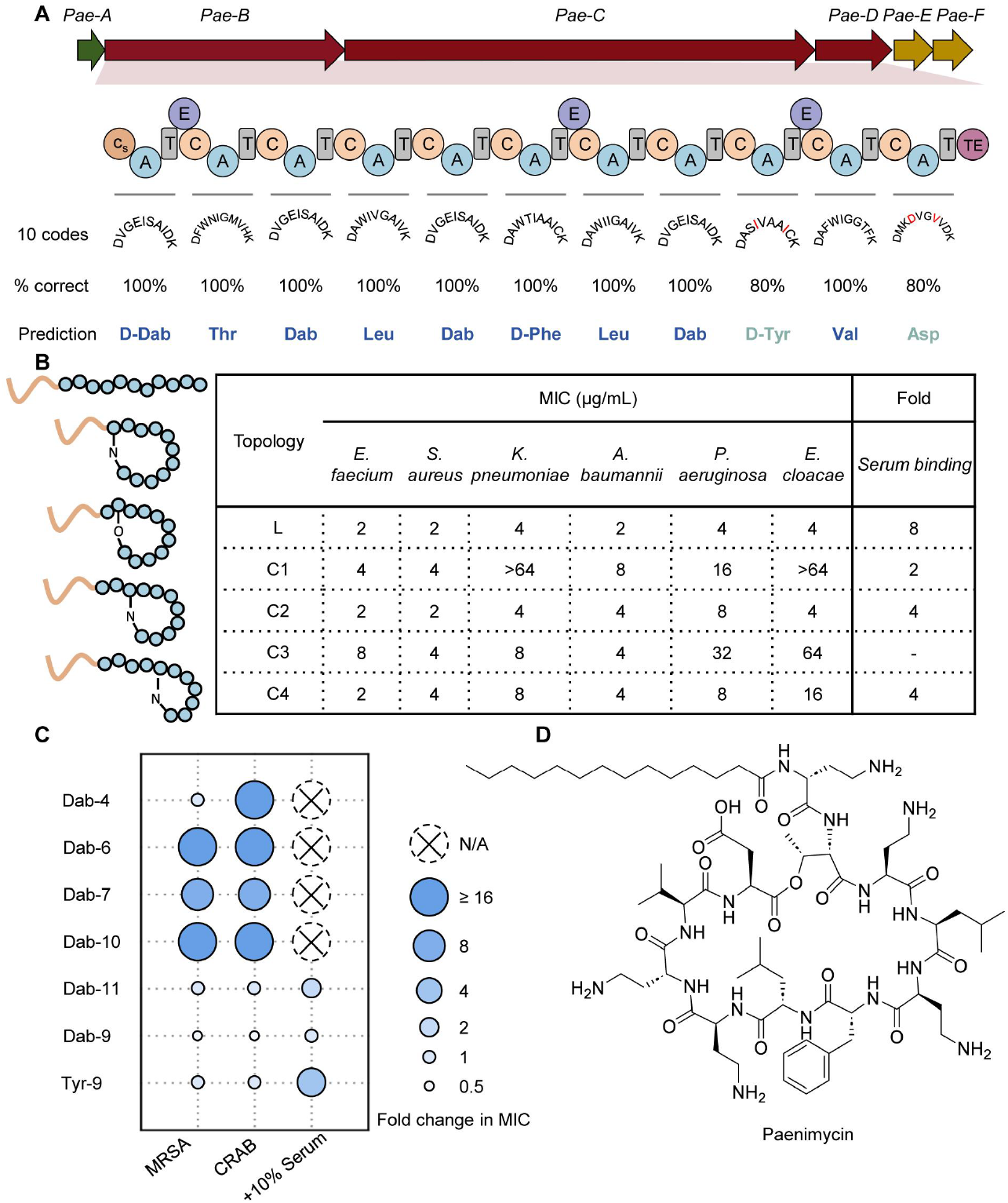
Structure optimization of BNP37 analogs leading to the discovery of paenimycin. A) The BGC of BNP37. 10-signature code sequences were used to predict each adenylation (A) domain, with the prediction accuracy was deemed reliable when it reached 80%. B) Diagrams of five peptide topologies predicted to be arisen from BNP37 BGC, along with their antimicrobial activity and serum binding efficacy against ESKAPE pathogens of all the peptides. (*E. faecium* BNCC 186301, *S. aureus* BNCC 186335, *K. pneumoniae* BNCC 186113, *A. baumannii* BNCC 194496, *P. aeruginosa* BNCC 186070, *E. cloacae* BNCC 185028). C) Fold changes in MIC following 2,4-Diaminobutyric Acid (Dab) screening. Specifically, all non-Dab positions were replaced with Dab respectively, MIC values against methicillin-resistant *S. aureus* (MRSA, *S. aureus* ATCC BAA44) and carbapenems-resistant *A. baumannii* (CRAB, *A. baumannii* ATCC BAA1605) with 10% serum were evaluated. N/A: not applicable. D) Structure of paenimycin.

### Antimicrobial spectrum

To comprehensively investigate the antimicrobial spectrum of paenimycin, we conducted microbroth susceptibility tests against a range of clinically prevalent MDR Gram-positive and Gram-negative pathogens, including the common hospitalized ESKAPE pathogens and WHO priority pathogens. Paenimycin exhibited broad-spectrum activity against all tested strains, with MIC values ranging from 2 to 4 μg/mL (Table 1). Notably, paenimycin demonstrated exceptional efficacy against Gram-negative bacteria, particularly those classified as critical and high priority by the WHO, such as carbapenem-resistant *Acinetobacter baumannii* (CRAB), third-generation cephalosporin-resistant or carbapenem-resistant *Enterobacterales*. While colistin is regarded as the last resort for treating MDR Gram-negative infections, its effectiveness is increasingly compromised by the global spread of the plasmid-borne mobilized colistin-resistance gene *mcr-1 (23)*. To date, 34 genetic variants and 10 alleles of mcr-1 have been identified worldwide, indicating its ongoing evolution and dissemination (*24, 25*). Paenimycin maintains potent activity (MIC=2 μg/mL) against colistin-resistant bacterial pathogens, including both acquired and intrinsic colistin-resistant strains. Furthermore, unlike colistin, paenimycin also shows comparable inhibitory activities against MDR Gram-positive bacteria, such as methicillin-resistant *Staphylococcus aureus* (MRSA) and erythromycin resistant *Streptococcus pyogenes* (ERSP), with a MIC value of 2 μg/mL. Importantly, paenimycin exhibits no activity against fungal pathogens at concentrations up to 64 μg/mL, indicating a low potential for cross-toxicity with mammalian cells.

**Table 1.** Antimicrobial activity of paenimycin against MDR bacteria.

|  | Pathogens | Strain Description | Resistant phenotype | MIC (µg/mL) |  |  |  |
| --- | --- | --- | --- | --- | --- | --- | --- |
|  |  |  |  | Paenimycin | Colistin | Ceftizoxime | Ciprofloxacin |
| Gram- | *K. pneumoniae* | ATCC 13883 | Amx<sup>R</sup>, Amp<sup>R</sup>, Cip<sup>R</sup> | 2 | 0.25 | 16 | 16 |
|  | *A. baumannii* | ATCC 19606 | Amx<sup>R</sup>, Amp<sup>R</sup>, Cip<sup>R</sup> | 2 | 0.25 | 16 | 32 |
|  | *A. baumannii* | BNCC 194496 | Cip<sup>R</sup> | 2 | 0.25 | 4 | 16 |
|  | *A. baumannii* | ATCC BAA1605 | Amx<sup>R</sup>, Amp<sup>R</sup>, Cza<sup>R</sup>,<br>Cip<sup>R</sup>, Gen<sup>R</sup>, Mer<sup>R</sup>, Ipm<sup>R</sup>,<br>Tcy<sup>R</sup> | 2 | 0.25 | >64 | >64 |
|  | *P. aeruginosa* | ATCC 27853 | Amx<sup>R</sup>, Amp<sup>R</sup>, Cip<sup>R</sup> | 2 | 0.5 | 16 | 2 |
|  | *E. coli* | ATCC 25922 |  | 2 | 0.25 | 0.125 | 2 |
|  | *E. coli* | clinical isolate 9 | Amx<sup>R</sup>, Amp<sup>R</sup>, Azm<sup>R</sup>,<br>Cza<sup>R</sup>, Cip<sup>R</sup>, Gen<sup>R</sup>, Ipm<sup>R</sup>,<br>Mer<sup>R</sup>, Tcy<sup>R</sup> | 2 | 0.25 | >64 | >64 |
|  | *E. coli* | clinical isolate 18 | Amx<sup>R</sup>, Amp<sup>R</sup>, Cza<sup>R</sup>,<br>Cip<sup>R</sup>, Gen<sup>R</sup>, Ipm<sup>R</sup>, Mer<sup>R</sup> | 2 | 0.25 | >64 | >64 |
|  | **Colistin acquired/intrinsic resistant** |  |  |  |  |  |  |
|  | *K. pneumoniae* | BNCC 359393 | Amx<sup>R</sup>, Amp<sup>R</sup>, Azm<sup>R</sup>,<br>Cza<sup>R</sup>, Cip<sup>R</sup>, Col<sup>R</sup>, Gen<sup>R</sup>,<br>Ipm<sup>R</sup>, Mer<sup>R</sup> | 2 | >64 | >64 | >64 |
|  | *E. cloacae* | ATCC 13047 | Amx<sup>R</sup>, Col<sup>R</sup>, Cza<sup>R</sup> | 2 | >64 | 8 | 4 |
|  | *E. cloacae* | BNCC 185928 | Amx<sup>R</sup>, Col<sup>R</sup>, Cza<sup>R</sup> | 4 | >64 | 8 | 2 |
|  | *A. baumannii* | BNCC194496-*mcr-1* | Col<sup>R</sup> | 4 | 16 | 4 | 16 |
|  | *E. coli* | MG1655-*mcr-1* | Col<sup>R</sup> | 2 | 32 | 0.0625 | 4 |
|  | *N. gonorrhoea* | BNCC 270346 | Col<sup>R</sup> | 2 | >64 | 1 | 16 |
|  | *N. meningitidis* | BNCC 353914 | Col<sup>R</sup> | 2 | >64 | 4 | 8 |
| Gram+ |  |  |  | Paenimycin | Colistin | Vancomycin | Methicillin |
|  | *S. aureus* | ATCC BAA44; | Met<sup>R</sup>, Amx<sup>R</sup>, Lin<sup>R</sup>, Ery<sup>R</sup>,<br>Azm<sup>R</sup>, Pen<sup>R</sup> | 2 | >64 | 1 | >64 |
|  | *S. aureus* | USA 300 | Met<sup>R</sup>, Amx<sup>R</sup>, Ery<sup>R</sup>, Pen<sup>R</sup>,<br>Azm<sup>R</sup> | 2 | >64 | 1 | >64 |
|  | *S. aureus* | BNCC 186335 | Azm<sup>R</sup>, Pen<sup>R</sup> | 2 | >64 | 0.5 | 1 |
|  | *S. pyogenes* | BNCC 336670 |  | 2 | >64 | 0.5 | 0.25 |
|  | *S. pyogenes* | 8007 | Ery<sup>R</sup> | 2 | >64 | 0.5 | 0.125 |
| Fungi | *Candida albicans* | BNCC 186382 | Fca<sup>R</sup> | >64 | >64 | >64 | >64 |
|  | *Aspergillus fumigatus* | BNCC 122691 | Fca<sup>R</sup>, 5-FU<sup>R</sup>, Tbf<sup>R</sup> | >64 | >64 | >64 | >64 |

Note: Amp<sup>R</sup>: Ampicillin resistance; Amx<sup>R</sup>: Amoxicillin resistance; Azm<sup>R</sup>: Azithromycin resistance; Cip<sup>R</sup>: Ciprofloxacin resistance Col<sup>R</sup>: Colistin resistance; Cza<sup>R</sup>: Ceftizoxime resistance; Ery<sup>R</sup>: Erythromycin resistance; FCA<sup>R</sup>: Fluconazole resistance; Gen<sup>R</sup>: Gentamycin resistance; Ipm<sup>R</sup>: Imipenem resistance; Mer<sup>R</sup>: Meropenem resistance; Lin<sup>R</sup>: Lincomycin resistance; Met<sup>R</sup>: Methicillin resistance; Pen<sup>R</sup>: Penicillin resistance; Tbf<sup>R</sup>: Terbinafine resistance; Tcy<sup>R</sup>: Tetracycline resistance; 5-FU<sup>R</sup>: 5-Fluorocytosine resistance.

### Antimicrobial mode of action

Given the notable activity of paenimycin against both MDR Gram-negative and Gram-positive bacteria, we next investigated its bactericidal mechanism. Through bacterial killing assays and scanning electron microscopic (SEM) analysis (fig. S5A-5C), we confirmed that paenimycin is a bactericidal antibiotic. At 1×MIC, it completely eradicated bacterial populations within 4 hours of incubation, causing cell collapse. The bacterial killing rate of paenimycin is faster than that of vancomycin while slower than that of colistin. Paenimycin did not induce cell depolarization but disrupted cell membrane integrity, leading to a dose-dependent release of potassium ions (fig S5D-5F). Given its broad-spectrum activity, these findings suggest that paenimycin likely targets the cell wall or cell membrane but employs a killing mechanism distinct from those of the narrow-spectrum antibiotics vancomycin and colistin. To identify the molecular target of paenimycin, we attempted to raise resistant mutants of *E. coli* ATCC 25922 and *S. aureus* BNCC 186335. Bacteria were exposed to serially increasing concentrations of paenimycin or maintained at sub-MIC levels of paenimycin for a continuous 30-day period (fig. S6). Despite testing under a variety of extreme conditions, we did not find any mutations resulting in an MIC increase greater than 2-fold. These findings, therefore, suggest that the target of paenimycin is most likely not proteins, as such targets are typically prone to mutations that can confer resistance. To further investigate, we screened the key bacterial cell components, including genomic DNA, total proteins and cell wall components such as peptidoglycan and lipopolysaccharide (LPS) (Fig 3A). Consistent with the findings from mutation assays, the addition of total genomic DNA and proteins did not affect the MIC values of paenimycin. However, LPS exhibited a dose-dependent inhibition of the antimicrobial activity of paenimycin against *E. coli*, with a 64-fold increase in the MIC value observed at a concentration of 0.5 mg/mL (Fig 3B). Furthermore, isothermal titration calorimetry (ITC) revealed a strong binding affinity between paneimycin and LPS, with an apparent dissociation constant (Kd) of 2.18 μM (Fig. 3C), suggesting that LPS is the primary target of paenimycin.

**Fig. 3.**
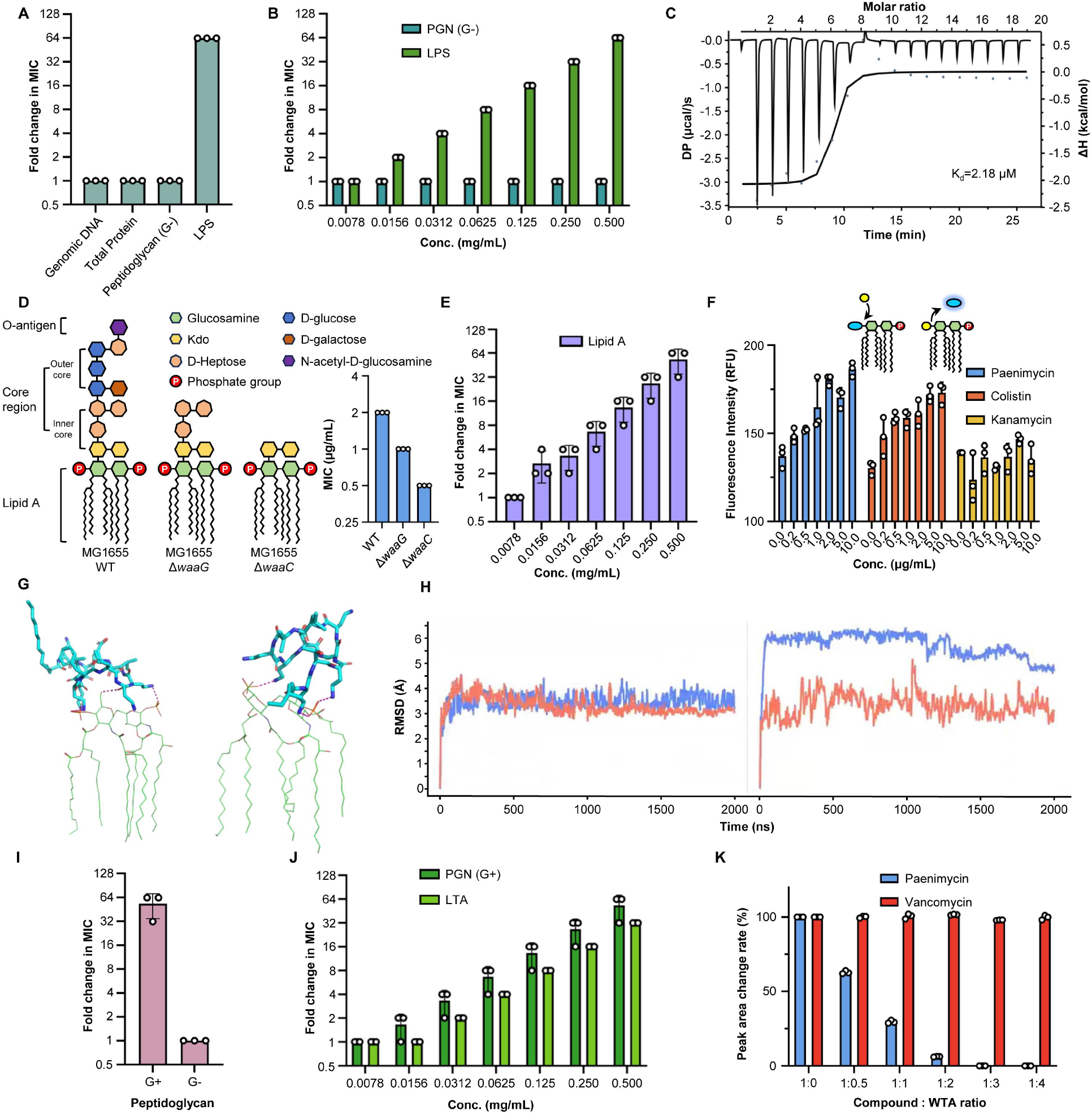
Target identification of paenimycin. A and B) Fold change in MIC of paenimycin against *E. coli* ATCC 25922 in the presence of genomic DNA (0.05 mg/mL) and total protein (0.38 mg/mL) extracted from the bacteria, alongside fixed concentration (0.5 mg/mL) and varying concentrations (0.0078-0.5 mg/mL) of peptidoglycan (PGN) and lipopolysaccharide (LPS). C) Isothermal titration calorimetry (ITC) assay determining the binding of paenimycin and LPS. D) Schematic chemical structures and MIC value changes of three different LPS molecules in *E. coli* MG1655 WT and gene knockout strains: *E. coli* MG1655 Δ *waaG* and *E. coli* MG1655 Δ *waaC*. E) Fold change in MIC of paenimycin against *E. coli* ATCC 25922 in the presence of different concentrations of lipid A (0.0078-0.5 mg/mL). F) Displacement of BODIPY TR fluorescence dye from lipid A binding upon addition of colistin, paenimycin, and kanamycin, with dye concentrations ranging from 0.0 to 10.0 mg/mL. G) Binding snapshots of paenimycin (left) and colistin (right) to lipid A from molecular dynamics (MD) simulations. Paenimycin and colistin are depicted as sticks, while lipid A is represented as a line; carbon atoms are colored cyan for paenimycin and green for colistin. Atomic interactions are shown as dashed purple lines. H) Root mean square deviation (RMSD) of MD simulation. The result for paenimycin was shown in left panel and colistin was shown in right panel. Lipid A was depicted in red and paenimycin or colistin was depicted in blue. I and J) Fold changes in MIC of paenimycin against *S. aureus* BNCC 186335 in the presence of fixed concentration (0.5 mg/mL) and varying concentrations (0.0078-0.5 mg/mL) of PGN of Gram-positive bacteria and lipoteichoic acid (LTA). K) Changes in the content of the remaining paenimycin and vancomycin in the solution after addition of wall teichoic acid (WTA) at different ratios.

LPS is a vital glycolipid and a key component of the outer membrane bilayer in Gram-negative bacteria. It consists of three structural domains: lipid A (a disaccharide backbone linked to multiple fatty acid chains anchoring LPS to the outer membrane), an core region (a branched oligosaccharide domain containing nine or ten sugars composed of an inner core and an outer core), an O-antigen (a repetitive carbohydrate polymer covalently attached to the core region) (*26, 27*) (Fig. 3D). To identify the specific LPS region that interacts with paenimycin, we knocked out the *waaC* and *waaG* genes (*28*). These genes, which are non-essential for bacterial survival, encode enzymes responsible for synthesizing the inner and outer cores of the LPS structure, respectively (fig. S7). Paenimycin maintained strong antimicrobial activity against the strains deficient in either the outer core or those lacking both the inner and outer cores (Fig. 3D), indicating that the inner and outer core regions of LPS are not binding sites for paenimycin. Next, to determine whether paenimycin binds lipid A, a crucial outer membrane component and primary target of colistin (*15*), we performed MIC assays of paenimycin against *E. coli* in the presence of varying concentrations of lipid A. Lipid A inhibited the antibacterial activity of paenimycin in a dose-dependent manner (Fig. 3E). Furthermore, paenimycin competitively displaced the lipid A-specific fluorescent dye BODIPY-TR, confirming its strong binding affinity for lipid A (Fig. 3F). Although both paenimycin and colistin share lipid A as a cellular target, paenimycin retains potent antibacterial activity against both intrinsic (e.g. EptA, PmrA/B and PhoP/Q genes) (*29*) and acquired (e.g. *mcr-1* gene) (*23*) colistin resistance strains, with a MIC value of 2 μg/mL (Table 1). Consistently, we observed that adding phosphoethanolamine (pEtN)-modified lipid A (31, 32), which is catalyzed by EptA and contributes to colistin resistance, reduced the activity of paenimycin in antibacterial assays, while it had no effect on colistin (fig S8). This suggests that paenimycin employs a novel mechanism of action, targeting a distinct site on lipid A to kill Gram-negative bacteria. To identify the specific binding site, we performed molecular dynamics simulations (Fig. 3G and 3H). The results revealed that paenimycin interacts with both the phosphate groups and notably the C-6 hydroxyl group of the glucosamine moieties. In contrast, colistin exclusively targets the phosphate groups. The additional interaction with the hydroxyl group likely account for paenimycin’s sustained efficacy against colistin-resistant strains.

Unlike colistin, paenimycin is a broad-spectrum antibiotic with potent activity against both Gram-negative bacteria and Gram-positive bacteria. Since lipid A, the target in Gram-negative bacteria, is absent in Gram-positive bacteria, this suggests that paenimycin may interact with an alternative target in Gram-positive bacteria. To investigate this issue, systematic feeding studies were conducted, revealing that peptidoglycan extracted from Gram-positive bacteria reduces the antibacterial activity of paenimycin against Gram-positive bacteria, whereas peptidoglycan from Gram-negative bacteria does not exhibit the same effect (Fig. 3I). The key difference between the peptidoglycan of Gram-positive bacteria and Gram-negative bacteria peptidoglycan is the presence of teichoic acids (TAs), including lipoteichoic acid (LTA) and wall teichoic acid (WTA) (*30, 31*). This led us to hypothesize that LTA or WTA might serve as the molecular target of paenimycin. To test this hypothesis, we conducted MIC assays of paenimycin with the addition of LTA. Results showed that LTA exhibited a dose-dependent inhibition of paenimycin’s activity, with 0.5 mg/mL completely abolishing its antibacterial effect (Fig. 3J). ITC analysis further confirmed a strong binding affinity between paenimycin and LTA, with an apparent Kd value of 5.61 μM (fig. S9A). Moreover, WTA also formed a complex with paenimycin, resulting in a measurable concentration decrease in the solution (Fig. 3K). Structurally, both LTA and WTA contain multiple phosphate groups, forming highly anionic substructures that may interact with the cationic regions of paenimycin. Molecular dynamics simulations further confirmed that the phosphate groups in LTA and WTA serve as the primary binding site of paenimycin (fig. S9B).

Our findings indicate that paenimycin binds to the phosphate and hydroxyl groups of lipid A of LPS in Gram-negative bacteria, as well as to the phosphate groups in LTA and WTA within the peptidoglycan of Gram-positive bacteria. This dual binding mechanism explains paenimycin’s broad-spectrum activity against MDR pathogens without detectable resistance (Fig. 4A). Given the broad-spectrum antimicrobial activity and attractive dual binding targets, we subsequently investigated the potential of paenimycin as an antimicrobial therapeutic agent. Paenimycin exhibited low toxicity against mammalian cells, with HC_50_ > 100 μg/mL and IC_50_ > 30 μg/mL against HepG2 cells, and LD_50_ >500 mg/kg (Fig. 4B, fig. S10A and 10B), suggesting paneimycin harbors a wide therapeutic window. Additionally, paenimycin showed favorable pharmacokinetic parameters using subcutaneous injection (s.c.), with the half-life time of 20.2±1.9 hours and bioavailability of 102±13% (Fig. 4C and table S5), which is 10 times higher than that of colistin. Notably, unlike the pharmacokinetic pattern of colistin, the half-life time of paenimycin administrated using s.c. is 2.4-fold longer than that administrated using intravenous (i.v.) injection (fig. S10D and table S5). Based on these findings, s.c. administration was selected as the primary route for *in vivo* studies of paenimycin. Given that nephrotoxicity is a primary concern associated with the use of antibiotics (i.e., colistin (*16, 17*), amphotericin B (*32*)), next generation of colistin analogues with decreased nephrotoxicity have been developed, including SPR-206 (phase I) (*33*), QPX-9003 (phase II) (*34*), and MRX-8 (phase I) (*35*). To evaluate the nephrotoxicity of paenimycin, mice were administrated with high concentration of paenimycin (i.e., 40 mg/kg) for a continuous 7 days. The kidney tissues were then analyzed using kidney injury biomarkers (including KIM-1, LCN2, SPP1 and TIMP-1) as well as histological evaluation, with colistin serving as a control antibiotic. Even at high concentration (20-fold of MIC) and prolonged exposure, paenimycin did not induce any detectable kidney injury. In contrast, colistin caused a severe nephrotoxicity, as indicated by both biomarkers and histological changes (Fig. 4D and fig. S10E).

**Fig. 4.**
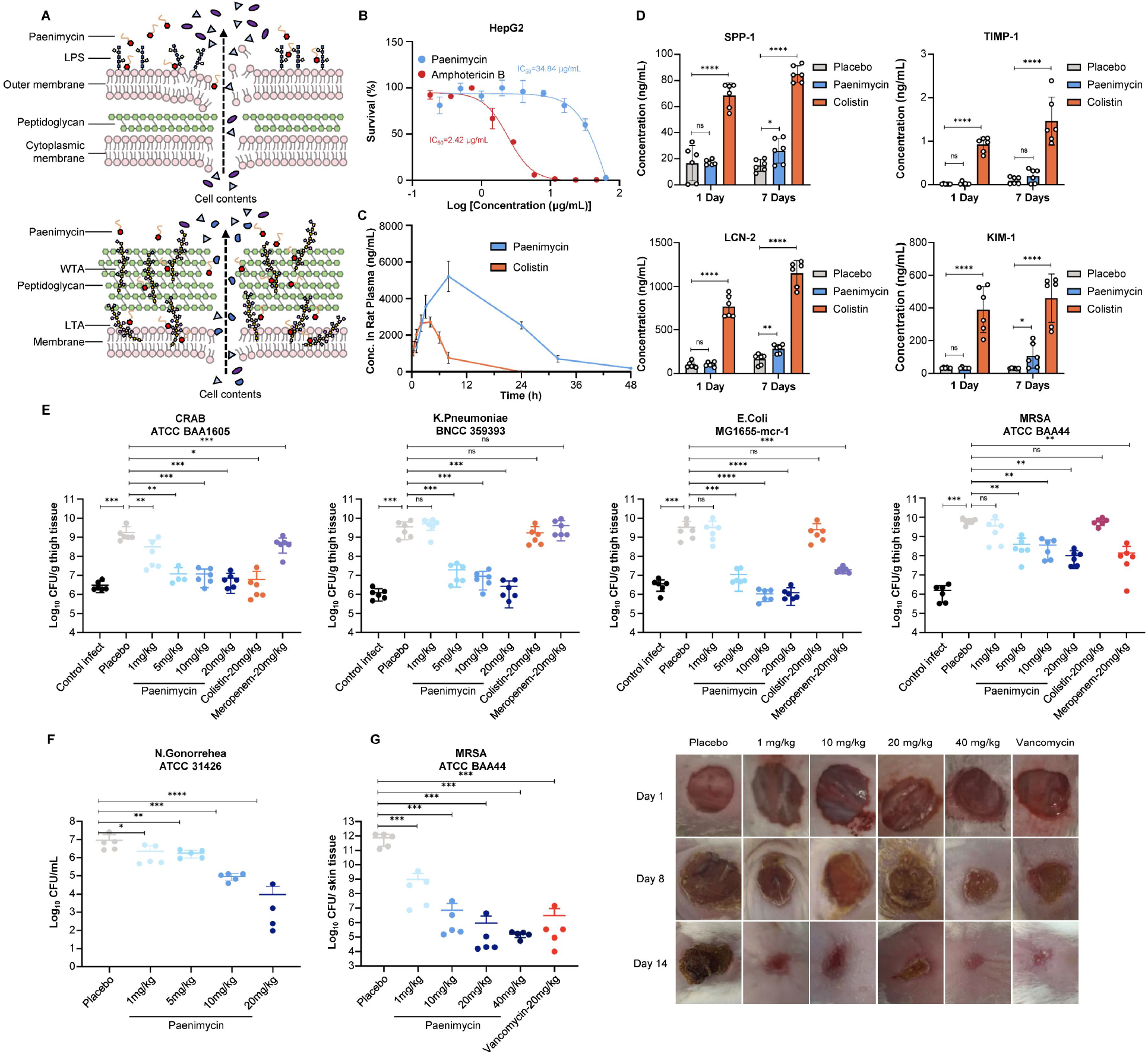
Toxicity and *in vivo* efficacy of paenimycin. A) Schematic models illustrating the proposed mode of action of paenimycin for Gram-negative (above) and Gram-positive (below) bacteria. B) Cytotoxicity assay of paenimycin against human hepatocellular carcinomas (HepG2) cell line. Amphotericin B was used as a toxic control. C) The pharmacokinetic curves of paenimycin and colistin over time after administration of 10 mg/kg via subcutaneous injection. D) *In vivo* nephrotoxicity assays of paenimycin and colistin. The concentrations of biomarkers (SPP1, TIMP-1, LCN2 and KIM-1) in mice serum were measured at 1 day and 7 days post administration of 40 mg/kg of paenimycin and colistin, respectively (s.c., qd, n=3 mice per group). E) Neutropenic thigh infection model using drug resistant strains (n=3 mice/ 6 thighs per group). All therapeutic compounds and vehicles were administrated via subcutaneous injection once daily and the bacteria burden were counted after 24 hours post drug treatment. Significant differences were analyzed by one-way analysis of variance (ANOVA) (\**p* ≤ 0.05, \*\**p* ≤ 0.01, \*\*\**p* ≤ 0.001, **** *p* ≤ 0.0001). F) Vaginitis infection model using N.gonorrehea ATCC 31426. All therapeutic compounds and vehicles were administrated via subcutaneous injection once daily for 3 consecutive days. On the day after the last administration, vaginal lavage fluid were collected and diluted for bacteria counting. Significant differences between each group with placebo were analyzed by t-test analysis (*p≤ 0.05, **p≤0.01, ****p≤0.0001). G) Skin infection model using MRSA ATCC BAA44. Paenimycin and vancomycin were administrated via subcutaneous injection once daily for 5 consecutive days and the bacteria burden of wound tissues were counted 15 days after infection (n= 5 mice per group). Photographic records were taken. Significant differences were analyzed by one-way analysis of variance (ANOVA) (\*\*\**p*≤ 0.001).

### *In vivo* efficacy in mice

Given the potent antimicrobial activity and favorable pharmacokinetic parameters, we next investigated the *in vivo* efficacy of paenimycin against MDR bacteria using three different types of infection models, including neutropenic thigh infection model, skin infection model, and vaginal infection model. Our initial evaluation focused on paenimycin’s efficacy against CRAB using neutropenic thigh infection mouse model. Paenimycin exhibited a dose-dependent reduction in bacterial burden. At a dosage of 20 mg/kg administered once daily, the bacterial burden decreased by 2.40 log10 compared to the vehicle control, which is comparable with that of colistin, demonstrating potent efficacy *in vivo* efficacy of paenimycin (Fig. 4E). Given paenimycin’s distinct antimicrobial mechanism and the increasing prevalence of colistin-resistant MDR pathogens, we further investigated its *in vivo* efficacy using mouse models infected with carbapenem and/or colistin-resistant XDR pathogens (Fig. 4E). At a dose of 20 mg/kg once daily, paenimycin reduced the bacterial burden by 3.42 log10 in colistin resistant *E. coli* and 3.12 log10 in colistin- and carbapenem-resistant XDR *K. penomoniae*, while colistin showed no activity to the strains tested (Fig. 4E). To evaluate the efficacy of paenimycin against intrinsic colistin-resistant pathogens, a vaginal infection model infected with colistin intrinsic-resistant *N. gonorrhoeae* was established (Fig. 4F). Paenimycin exhibited a dose-dependent reduction, achieving a reduction of 2.83 log10 at a dose of 20 mg/kg per day via s.c. administration. Furthermore, in the treatment of MDR Gram-positive bacteria, paenimycin showed dose-dependent activity against MRSA in both the neutropenic thigh infection model and the wound infection model (Fig. 4E and 4G). At 20 mg/kg once daily via s.c. administration, paenimycin reduced the bacterial burden by 1.79 log10 and 5.90 log10, respectively, comparable to the reductions achieved with vancomycin. The low nephrotoxicity, favorable pharmacokinetic profile, and potent *in vivo* efficacy collectively suggest that paenimycin is a promising drug candidate for treating difficult-to-manage infections encountered in clinical settings, particularly those caused by colistin-resistant MDR pathogens.

## Conclusion

Infections caused by MDR bacterial pathogens represent a critical threat to global public health. The growing prevalence of MDR pathogens, coupled with the declining rate of novel antibiotic discovery, underscores the urgent need for innovative therapeutic strategies. *Paenibacillaceae* are a rich source of antimicrobial natural products and leveraging these natural resources is expected to offer new solutions for combating resistant infections. Applying the culture-independent synBNP approach combined with activity-guided screening, we synthesized and screened 74 peptides whose structures were predicted based on the adenylation domain specificity of *Paenibacillaceae* NRP BGCs. This effort led to the discovery of a novel class of BNP37 peptides exhibiting potent, broad-spectrum inhibitory activity against MDR pathogens. Subsequent structural optimization of the BNP37 peptides led to the identification of paenimycin, an 11-mer depsi-lipopeptide featuring five positively charged Dab moieties and an N-acylated myristic acid chain, and with enhanced broad-spectrum antibacterial activity and reduced serum-binding affinity. Mechanistic studies revealed that paenimycin interacts not only with the phosphate groups of lipid A but also uniquely targets the hydroxyl group of glucosamine in lipid A, a key interaction enabling its potency against both intrinsic and acquired colistin-resistant pathogens in both in vitro and in vivo settings. Furthermore, paenimycin’s ability to bind the phosphate groups of teichoic acids in Gram-positive bacteria contributes to its broad-spectrum efficacy. Its unique mechanism of action, broad-spectrum activity, absence of detectable resistance, favorable pharmacokinetic profile, low nephrotoxicity, and excellent *in vivo* efficacy collectively position paenimycin as a promising candidate for treating infections caused by MDR pathogens, particularly those resistant to carbapenems, methicillin, and colistin. Due to the limited genomic data available for *Paenibacillaceae* bacteria, our study does not encompass all the types of potential NRPs encoded by this family. Furthermore, the structures of peptides derived from some NRP BGCs cannot be accurately predicted due to the presence of one or more unusual amino acids that current algorithms are unable to reliably forecast. Future work will focus on expanding the genomic data for *Paenibacillaceae* bacteria and advancing computational methods to more accurately predict unusual amino acids in NRPs, both of which are expected to enhance efforts in discovering new antibiotics to combat MDR infections.

## Author contributions

Z.Q.W conceived the project; Z.Q.W, W.Y.H, X.T.H contributed to manuscript writing, review, and editing; Z.Q.W, W.Y.H contributed to data analysis. W.Y.H performed the biochemical experiments. X.T.H performed the peptide synthesis. Q.S.D contributed to the ITC experiments and nephrotoxicity tests. Y.C.L contributed to the genetic knockout. T.C performed the data collection and bioinformatic analysis. W.X contributed to the MD simulation. W.Y.H, X.T.H and L. M. M contributed to the animal study. Y.J.C, Z. S, and N.L contributed to manuscript review. All authors contributed to data analysis and interpretation.

## Data and materials availability

The genomic sequence of *Paenibacillus caseinilyticus* GW78 is available in GenBank under accession number NZ_JAQAGY010000015.1. HPLC, HRMS and NMR spectra for paenimycin are presented as Supplementary Information. All data are available in the main text or the supplementary materials. All data, code, and materials used in the analysis must be available in some form to any researcher for purposes of reproducing or extending the analysis.

