## Supplementary data for "A novel broad-spectrum antibiotic targets multiple-drug-resistant bacteria with dual binding targets and no detectable resistance"

**Affiliations:**

**The PDF file includes:**

Materials and Methods
Figs. S1 to S16
Tables S1 to S6
References

**Other Supplementary Materials for this manuscript include the following:**

### Materials and Methods

#### Identification and bioinformatic analysis of the paenimycin (*pae*) BGC:

To comprehensively explored the non-ribosomal peptides encoded in the family of paenibacillaceae bacteria, all assembled sequences from publicly available genomic database which encompassed 1245 genomes, as well as 11 sequencing data derived from the sequencing of collected strains were retrieved. The Raw sequenced reads were assembled into contigs using SPAdes (version 3.13.0), with default parameters. Contigs were refined using Ragtag (version 2.1. 0) to obtain scaffold-level genomic data. A total of 17037 BGCs were retrieved using antiSMSH (version 7.0. 0) software analysis for all genomic data. An in-house Python script using a json module from Python was written to filter BGCs. In this analysis, the Python script used to read each JSON file produced by antiSMASH one at a time, gathering information on the kind of BGC, the quantity of A domains, and the presence or absence of a TE domain. We focused exclusively on NRPS BGCs containing a TE domain and 5 to 15 A domains for our study. The resulting 879 BGCs were further analyzed by clustering their GenBank files using Big-Scape (version 1.1.5), and the Clustering results were visualized using Cytoscape (version 3.10.0).

#### Peptide synthesis:

All candidate peptides were synthesized using standard solid-phase peptide synthesis (SPPS) methods on 2-chlorotrityl chloride resin. 2-chlorotrityl chloride resin (0.3g) was swollen in dichloromethane (DCM) for 30 minutes at room temperature. The first Fmoc-protected amino acid (3 equiv.) was loaded using 0.3 mL 2,4,6-collidine in 8 mL DCM for 10 hours, and then capping the unreacted Cl groups with 10 mL mixed reagents (Methanol: N,N-Diisopropylethylamine (DIPEA) : DCM = 1 : 0.5 : 9) for 45 minutes at room temperature. Fmoc deprotection was carried out using 20% piperidine in *N*,*N*-Dimethylformamide (DMF) (3 mL) for 7.5 minutes (2X) and then the resin was washed thoroughly with DMF (3 mL, 5X). Coupling reactions were carried out using Fmoc-protected amino acids (3 equiv.) pre-mixed with O-(7-Azabenzotriazollyl)-N,N,N',N'-tetramethyluronium hexafluophosphate (HBTU) (3 equiv.) and DIPEA) (3 equiv.) in DMF. Each coupling reaction was carried out for at least 1 hour and then the resin was washed with DMF (3 mL, 5X). The coupling steps were repeated as described above until the last amino acid (or a lipid acid) in the sequence was loaded.

1) Amide bond formation: Amide bonds were formed between the α-amino group of the N-terminal amino acid or amino acids with an animo group on the side chain and α-carboxyl group of C-terminal amino acid. Alloc-protected animo acid was used to selectively form the amide bond at the desired position. The resin-bound linear peptide was washed with DCM (3 mL, 5X) and treated with phenylsilane (15 equiv.) and tetrakis (triphenylphosphine) palladium (0) (0.5 equiv.) in DCM and shake 2h at room temperature, followed by washing with 10% sodium diethyldithiocarbamate trihydare in DMF (50 mL) and DCM (6 mL, 5X).

2) Peptide cyclization: Linear peptides were cleaved from resin by treating with 1% Trifluoroacetic acid (TFA) in DCM for 5 minutes and collected in a falcon tube. Then the cleaved peptides were mixed with benzotriazol-1-yl-oxytripyrrolidino-phosphonium hexafluorophosphate (PyBOP) (8 equiv.) and DIPEA (30 equiv.) in 45 mL DCM and shaken overnight.

3) Ester bond formation: Ester bonds were formed between the unprotected hydroxyl group of serine or threonine and the α-carboxyl group of the C-terminal animo acid. The resin-bound

peptides were mixed with amino acid (20 equiv.), DIPEA (40 equiv.), benzoyl chloride (20 equiv.) and 4-(dimethylamino)-pyridine (0.8 equiv.) in 15 mL DCM and rock for 48 hours.

4) Final cleavage: 5 mL of cleavage cocktail (95% TFA, 2.5% triisopropylsilane and 2.5% water) was added to the air-dried peptide mixture and shaken for 2 hours. Then the solution was added to a cold mixture of diethyl ether:hexane (1:1) and kept at -20 ℃ for 1 hour to precipitate the crude peptides.

**Minimum inhibitory concentration (MIC) assay:**
MIC assays were conducted following the protocol recommended by the Clinical and Laboratory Standards Institute (CLSI). Culture conditions are detailed in supplementary table S4. All synthetic compounds were dissolved in sterile dimethyl sulfoxide (DMSO) (SCR, CN) to a stock concentration of 12.8 mg/ml. Colistin, ceftizoxime and ciprofloxacin were used as positive controls for Gram-negative bacteria, while vancomycin and methicillin were used as positive controls for Gram-positive bacteria. Compound stock solutions were serially diluted across 96-well plates using a 2-fold dilution to give a concentration up to 64 µg/ml in a volume of 50 µL per well. Overnight cultures of each assayed strains were diluted 5000-fold in fresh medium and 50 µL of those dilutions were added into each well. The lowest concentration with no visible growth of bacteria after 16 hours was recorded as the MIC value. All assays were performed in duplicate and repeated three independent times.

**Cell lysis assay:**
A single colony of *S. aureus* BNCC 186335 was inoculated overnight at 37 ℃ with shaking at 220 rpm. Cells were collected, resuspended in sterile PBS (pH 7.4) and then diluted to an $OD_{600nm}$ of 0.35. 900 µL of this bacteria suspension was mixed with 100 µL of 17 µM SYTOX green nucleic acid stain (Thermo Fisher, USA). The mixture was incubated in the dark at 37 ℃ for 5 minutes, and then transferred to a 384-well flat bottom black microtiter plate (30 µL per well). The initial fluorescence intensity of each well was measured using a microplate reader (Infinite 200 Pro, Tecan, USA) at 9 second time intervals for 5 minutes (Excitation/Emission = 488/523 nm). The same volume of paenimycin solutions at concentration of 1, 2, 4, 8, 10×MIC were added into each well, and melittin and DMSO were used as positive and negative controls, respectively. The fluorescence intensity was then continually monitored for 25 minutes and plotted by Prism 9.0. Experiments were performed three independent times (n=3).

**Membrane depolarization assay:**
A single colony of *S. aureus* BNCC 186335 was cultured overnight at 37 ℃ with shaking at 220 rpm. Cells were harvested, washed in 5mM HEPES buffer, and resuspended in 5mM HEPES buffer with 20mM glucose ($OD_{600nm}$ = 0.1). 2 µL of 500 µM 3,3'-Dipropylthiadicarbocyanine Iodide [DiSC3(5)] dye (Macklin, CN) was added to 1 mL of the cell suspension and incubated in the dark at 37 ℃ for 30 minutes. 100 µL of this mixture was then added to a 384-well flat bottom black microtiter plate. The initial fluorescence intensity of each well was recorded using an Spectra Max ID5 (Molecular devices, USA) (Excitation/Emission = 620/670 nm) at 15 second time intervals for 5 minutes. 1 µL of paenimycin solutions at concentration of 1, 2, 4, 8, 10× MIC were added into each well, and Triton X-100 and DMSO were used as positive and negative controls, respectively. The fluorescence intensity was continually monitored for 20 minutes and plotted by Prism 9.0. Experiments were performed three independent times (n=3).

**Potassium ion release assay:**
The release of potassium ion was monitored using an Orion Dual Star pH/ISE Benchtop Meter (Thermo Fisher, USA). A single colony of *S. aureus* BNCC 186335 was inoculated and incubated overnight in 50mL LB medium at 37 ℃ with shaking at 220 rpm. Cells were harvested, washed twice with buffer (10 mM Tris-acetate, 100 mM NaCl, pH 7.4). and then resuspended in the same buffer and adjusted to an OD600nm of 1.0. The concentration of extracellular potassium ions was monitored at 5 second intervals. Paenimycin at concentrations of 1, 2, 4, 8×MIC and gramicidin at 8×MIC were added into the suspension respectively after 60 seconds and the measurement was continued until 8 minutes. Data were plotted by Prism 9.0. The experiments were performed in triplicate (n=3).

**Time-dependent killing assay:**
An overnight culture of *S. aureus* BNCC 186335 and *E. coli* ATCC 25922 was prepared by inoculating and incubating a single colony at 37 ℃ with shaking at 220 rpm, respectively. The cultures were diluted to a final concentration of $1\times10^6$ CFU/mL. 4 mL bacteria dilutions were then challenged with 1, 4 and 8×MIC of paenimycin, 8× MIC of vancomycin or 8× MIC of colistin. After 1, 2, 4, 8 and 16 hours, 100 µL aliquots of each culture were collected, serially 10-fold diluted and plated on LB agar plates. As for *E. coli*, additional time intervals at 0.25, 0.5, 0.75 hours were added. Colonies were counted after overnight growth at 37 ℃. The experiments were performed in three biological replicates (n=3).

**Scanning electron microscopy:**
Overnight cultures of *S. aureus* BNCC 186335 and *E. coli* ATCC 25922 were diluted 1:200 in fresh medium and grown to $OD_{600nm}$ of 0.8. Bacteria were treated with 8×MIC of paenimycin for 0, 1 and 4 hours, respectively. Cells were harvested and washed twice with PBS buffer (pH 7.4), and then fixed with 2.5% glutaric dialdehyde solution at 4 ℃ for 12 hours and rinsed three times with PBS buffer. The samples were postfixed with 1% osmium tetroxide solution for 2 hours at room temperature. After removing the solution, the samples were rinsed with PBS buffer three times. The samples were dehydrated in a graded series of ethanol concentrations (30%, 50%, 70%, 80%, 90%, 95%) for 15 minutes each and two times in 100% ethanol for 20 minutes each and put into a critical point dryer (Quorom k850, UK) to dry out. Subsequently, the samples were fixed on the sample stage using conductive carbon glue and sprayed Pt with the ion type sputtering instrument (Hitachi MC1000, JP) for 120 seconds. Imaging was carried out in a Hitachi Regulus8100 at 3.0 kv.

**Serial passaging assay:**
A single colony of *S. aureus* BNCC 186335 and *E. coli* ATCC 25922 was inoculated and incubated at 37 ℃ with shaking at 220 rpm overnight, respectively. The overnight cultures were diluted 1:5,000 into fresh medium and the MIC values of paenimycin, ciprofloxacin and bacitracin for *S. aureus* and paenimycin, ciprofloxacin and tetracycline for *E. coli* were tested using the MIC method and the MIC values were recorded after 24 hours. For bacitracin, the LB medium was supplemented with 50 µg/mL $ZnCl_2$. On the next day, the bacteria cultures of the sub-MIC well of each antibiotic were diluted into fresh medium in a 1:500 ratio and mixed with serial diluted antibiotics. Then the new MICs were tested following the method described above. This serial passage process lasted for 28 days. The experiments were conducted in triplicates.

**Feeding assay:**
The effect of bacteria cell components on paenimycin's antibacterial activity was evaluated using *S. aureus* BNCC 186335 and *E. coli* ATCC 25922. Cell components of Gram-positive and Gram-negative bacteria including peptidoglycan, total protein, genomic DNA, LPS and LTA were dissolved in water to a stock concentration of 5 mg/mL. Lipid A was dissolved in chloroform and premixed with paenimycin for 15 minutes and then the premixed solutions were evaporated by air. Each solution was added to individual wells of 96-well plates, and serially diluted to final concentrations of 0.0078 to 0.5 mg/mL. The MIC values were evaluated using the same method described for the MIC assay. Fold changes in MIC were calculated based on the formula: Fold change in MIC = Final MIC/Original MIC. The experiments were conducted in duplicate and repeated three independent times.

**Isothermal calorimetry (ITC) assay:**
The binding of paenimycin and LPS and LTA were measured using ITC. The ITC experiments were performed on PEAQ-ITC (Malvern, UK) instrument at 25 °C using a solution of 5 mM of paenimycin or 1 mM of colistin along with 100 μM LPS or LTA in 5 mM HEPES buffer (pH 7.4). The titration process involved an initial injection of 0.23 μL, followed by 18 injections of 2 μL at 80-second intervals, with continuous stirring at 500 rpm. Data were analyzed using PEAQ-ITC software, and the thermodynamic parameters [enthalpy (ΔH), entropy (ΔS), and the equilibrium binding constant (Kd)] were calculated using a one-binding site model.

**Knockout of *WaaC* and *WaaG* in Escherichia coli MG1655:**
Genome editing was conducted as described previously(*1*). The N20 sequence was inserted into ptargetF by Gibson assembly yielding the plasmids pTargetT-Δ*WaaC* and pTargetT-Δ*WaaG*. two fragments, one upstream and one downstream of the targeted gene, were PCR-amplified from the genomic DNA of Escherichia coli MG1655 using primer pairs *WaaC*_KOUF/ WaaC_KOUR and *WaaC* _KODF/ *WaaC* _KODR for *WaaC* and *WaaG*_KOUF/ *WaaG* _KOUR and *WaaG* _KODF/ *WaaG* _KODR for *WaaG*, respectively. Each pair of PCR products was constructed into a PCR fragment by overlapping PCR, yielding the donor DNA. *E. coli* MG1655 competent cells were prepared for harboring pCas. Arabinose (10mM final concentration) was added to the culture for λ-Red induction. For electroporation, 50 μL of cells was mixed with 100 ng of pTargetT series DNA and 400 ng of donor DNA. Cells were recovered at 30°C for 1 h before being spread onto LB agar containing kanamycin (50 mg/L) and spectinomycin (50 mg/L) and incubated overnight at 30 °C. Transformants were identified by colony PCR and DNA sequencing. For the curing of pTarget series DNA, the edited colony harboring both pCas and pTarget series was inoculated into 2 ml of LB medium containing kanamycin (50 mg/L) and IPTG (isopropyl-β-D-thiogalactopyranoside; 0.5 mM). The culture was incubated for 8 to 16 h, diluted, and spread onto LB plates containing kanamycin (50 mg/L). The colonies were confirmed as cured by determining their sensitivity to spectinomycin (50 mg/L). The colonies cured of pTarget series DNA were used in a second round of genome editing. pCas was cured by growing the colonies overnight at 37°C non selectively.

**BC displacement assay:**
The affinity of paenimycin to lipid A was assessed by evaluating its ability to displace BODIPY TR cadaverine (BC) from lipid A-BC mixture. 20 μg/mL LPS and 20 μM BC were premixed in 5 mM HEPES (pH7.4) and incubated at 37 ℃ for 30 minutes. Subsequently, the mixture was

transferred into a 96-well flat bottom black microtiter plate (50 µL per well). 50 µL of paenimycin in 5 mM HEPES were added at final concentrations ranging from 0.0 µg/mL to 1.0 µg/mL. Colistin and kanamycin were used as positive and negative controls, respectively. After incubating in the dark for 5 minutes, the fluorescence intensity was measured using a microplate reader (Infinite 200 Pro,Tecan) (Excitation/Emission = 580/620 nm) and plotted by Prism 9.0.

**Lipid A extraction and feeding assay:**
Lipid A and pEtN modified lipid A were extracted from *E. coli* MG1655 and *E. coli* MG1655-*mcr-1* following the previously described method(*2, 3*). Briefly, chemical lysis of bacterial cells was performed using a mixture of chloroform, methanol and water (Bligh-Dyer) solvent, and the LPS was pelleted by centrifugation. Then a combination of mild-acid hydrolysis and solvent extractions were used to extract lipid A from the pellet mixture. Lipid A can be defined as the chloroform soluble portion of LPS after mild-acid hydrolysis. The lyophilized crude lipid A was dissolved in chloroform and added into 96-well plates containing 4×MIC of paenimycin or colistin and diluted *E. coli* MG1655 solution following the feeding assay method. Then the $OD_{600nm}$ was measured using Multiscan SkyHigh at 37 ℃ for 16 hours and plotted using Prism 9.0. The experiments were performed in triplicates (n=3).

**Molecular Dynamic (MD) Simulations:**
Lipid A and colistin structures were extracted from PDB 1QFF and 8DEV, respectively(*4, 5*). Lipoteichoic acid (LTA) and paenimycin 3D model was optimized from 2D structure using MOE. Initial complex model was obtained by placing ligand close to receptor and fulfills the ionic charge interactions between Receptor (Lipid A or LTA) and Ligand (Colistin or paenimycin). The complex model was then sent for MD simulation with Desmond (Desmond/Maestro noncommercial version 2022.1). Specifically, 1,2-dipalmitoyl-phosphatidylcholine (DPPC) membrane model was first added automatically, then the DPPC orientation was modified to accommodate hydrophobic tail of the ligand to the center of membrane plane. Complexes were neutralized with Na+ or Cl- ions, and solvation was conducted with a 10 Å orthorhombic TIP3P water buffer box containing 0.15 M NaCl using default parameters in Desmond. MD simulation includes a 5-step minimization with restraints released gradually, and a 200 ns production run without restraints in NPgT ensemble. The first step of the minimization is to restrain solute heavy atoms at 10 K in NVT ensemble for 100 ps. The second is to restrain membrane in z-axis and protein atoms at 100 K in NPT ensemble for 20 ps. The third is to restrain membrane in z-axis and protein atoms at 100 K in NPgT ensemble for 100 ps. The fourth is to heat up from 100 to 300 K in NPgT ensemble for 150 ps. The fifth is removing all restraints in NVT ensemble for 100 ps. The 200 ns production simulation was conducted at 300 K under 1 bar in NPgT ensemble, and repeated 3 times, trajectory coordinates were saved every 100 ps.

**Wall teichoic acid extraction and binding assay:**
Wall teichoic acid was extracted from *S. aureus* BNCC 186335 as previously described(*6*). Briefly, wall teichoic acid was isolated from the crude peptidoglycan sacculus after completely washing with 4% SDS to remove lipoteichoic acid. Then hydrolysis with trichloroacetic acid was used to liberate water soluble WTA from peptidoglycan. The lyophilized crude WTA was dissolved in water and diluted to 1 mg/mL, 2 mg/mL, 3 mg/mL, 4 mg/mL and 5 mg/mL and then

mixed with 1 mg/mL paenimycin for 0:1, 1:1, 2:1, 3:1, 4:1 and 5:1 w/w ratios, respectively. Mixtures were incubated for 1 hour at room temperature and centrifuged for 5 minutes at 15,000 g to pellet insoluble material. The content of paenimycin in the supernatant was detected by UPLC (Waters, US). Peak aera change rate was calculated and plotted by Prim 9.0.

**Cytotoxicity assay:**
The cytotoxicity of paenimycin and its analogs was tested using the 3-(4,5-dimethyl-2-thiazolyl)-2,5-diphenyl-2H-tetrazolium bromide (MTT) assay. HepG2 cells were cultured in Dulbecco's Modified Eagle Medium (DMEM) supplemented with 10% fetal bovine serum. Cells were seeded into a 96-well plate with a density of 5,000 cells per well and cultured at 37 ℃ with 5% $CO_2$ for 24 hours. Then the medium was aspirated and replaced with 100 μL fresh medium containing compounds at a final concentration ranging from 64 μg/mL to 0.25 μg/mL, and amphotericin B was used as control. After 48 hours of incubation, the medium was removed and 110 μL of freshly prepared MTT solution (0.5 mg/mL) was added to each well. The plate was incubated for another 3 hours at 37 ℃ to allow crystal formation. Subsequently, precipitated formazan crystals were dissolved by addition of 100 μL of solubilization solution (40% DMF, 16% SDS and 2% acetic acid in H2O). The absorbance of each well was measured at OD570nm using a microplate reader (Multiscan SkyHigh, Thermo Fisher). $IC_{50}$ values were calculated using Prism 9.0 as the concentration of each compound required to inhibit 50% of cell growth. All the experiments were performed in triplicates (n=3) and repeated two independent times.

**Hemolytic assay:**
Fresh sterile defibrated sheep blood was centrifuged and resuspended in PBS (pH 7.4) and diluted to a concentration of $1\times10^9$ cells/mL. Serial dilutions of Paenimycin were prepared at concentrations ranging from 100 μM to 0.38 μM and mixed with the blood cell suspensions to yield a final volume of 500 μL. 1% Triton X-100 and 1% DMSO were used as positive and negative controls respectively. After a 3-hour incubation time at 37 ℃, the mixture was centrifuged at 3,000 rpm for 20 minutes and the supernatant was collected and transferred into a 96-well plate. The hemolysis efficacy was determined by measuring the absorbance at $OD_{540nm}$ using Multiscan SkyHigh and calculated using Prism 9.0. All the experiments were performed in biological triplicates (n=3).

**Pharmacokinetic study:**
All studies were approved by Pharmaron's Institutional Animal Care and Use Committee (IACUC). Six to eight-week-old SD male rats (200-300 g) were used in pharmacokinetic studies. Paenimycin was administered via intravenous (IV, 5 mg/kg) and subcutaneous (SC, 10 mg/kg) injection. Blood timepoints were taken after dosing at pre, 5 minutes, 15 minutes, 30 minutes, 1 hour, 2 hours, 4 hours, 6 hours, 8 hours, 24 hours and 48 hours for IV dosing, and pre, 1 hour, 3 hours, 8 hours, 24 hours, 32 hours, 48 hours, 56 hours, 72 hours and 96 hours for SC (10 mg/kg) dosing. Aliquots of 0.1 mL of blood were taken by puncture of the jugular vein from each rat (n = 3 per route and dose) at each timepoint. Samples were transferred into plastic micro centrifuge tubes with EDTA-$K_2$ and stored on ice. The blood samples were centrifuged at 4000g for 5 minutes at 4°C to obtain plasma within 30 minutes. The plasma samples were stored at -80 °C for further analysis.
HPLC-MS/MS pharmacokinetic analysis: paenimycin (l mg/mL, in DMSO) was serially diluted in ACN: water (50:50) to prepare the standard curves and quality control (QC) spiking solutions.

Standards and QCs were created by adding 5 μL of spiking solutions to 50 μL of drug free plasma (SD $K_2EDTA$ Rat, Vital River). 55 μL of standards, 55 μL of QC samples and 55 μL of unknown samples (50 μL of rat plasma with 5 μL of blank solution) were added to 200 μL of acetonitrile containing 10ng/mL of the internal standard (IS) Dexamethasone (Shanghai Aladdin Biochemical Technology Co.,Ltd) for precipitating protein respectively.
Extracts were vortexed for 1 minute and centrifuged at 4000 rpm for 15 minutes. The supernatant was diluted 3 times with water. 20 μL of diluted supernatant was injected into the LC/MS/MS system for quantitative analysis. HPLC-MS/MS analysis was performed on a Sciex Applied Biosystems Triple Quad 5500+ or 6500+ mass spectrometer coupled to a Shimadzu Nexera Series System Controller CBM-40 UPLC (Ultra-High- Performance Liquid Chromatography) system to quantify each drug in the plasma. Chromatography was performed on a Raptor Biphenyl column (3x30 mm; particle size, 2.7 μm) using a reversed phase gradient. 5% acetonitrile in ultrapure water with 0.1% formic acid was used for the aqueous mobile phase and 95% acetonitrile in ultrapure water with 0.1% formic acid was used for the organic mobile phase. Multiple-reaction monitoring (MRM) of parent/daughter transitions in electrospray positive-ionization mode was used to quantify the analytes. The following MRM transitions were used for paenimycin (467.00/461.50) and dexamethasone (393.06/373.00). Sample analysis was accepted if the concentrations of the quality control samples were within 20% of the nominal concentration. Data processing was performed using Analyst software (v1.7.3; Applied Biosystems Sciex).

**Nephrotoxicity assay:**
Pathogen-free female ICR mice (Hangzhou Medical College, China), aged six weeks and weighing between 23-27 g, were used in the study. The mice were randomly grouped into six individual cages with three mice per group. Mice were acclimatized for three days prior to the experiments. Paenimycin was formulated in a solution containing 10% DMSO and 0.5% Tween 80 in 0.9% saline. Colistin or a solvent without any antibiotics was used as the positive control and placebo, respectively. The compounds were administered by subcutaneous injection at a dosage of 40 mg/kg once daily for 1 day or 7 consecutive days. Serum was collected from blood samples 24 hours after the last dose. The concentration of nephrotoxicity-related biomarkers including kidney injury molecule-1 (KIM-1), tissue inhibitor of metalloproteinase-1 (TIMP-1), lipocalin-2 (LCN-2), and secreted phosphoprotein 1 (SPP-1) were measured using the commercial kits (Cloud-Clone Corp, China) and the ELISA assays were conducted following the manufacturer's instructions. All animals were euthanized, and the kidney tissues were collected, fixed, dissected, and subjected to H&E staining.

**Neutropenic thigh infection model:**
Pathogen-free female ICR mice (Hangzhou Medical College, China), aged six weeks and weighing between 23-27 g, were used in this study. The mice were housed in individually ventilated cages and acclimatized for three days before the experiments. Then the mice were rendered neutropenic via intraperitoneal injections with 150mg/kg and 100mg/kg of cyclophosphamide on day -4 and day -1, respectively. On day 0, mice were infected by intramuscular administration of 50 μL of *S. aureus* ATCC BAA-44, *E. Coli* MG1655-*mcr-1*, *A. baumannii* ATCC BAA-1605 and *K. pneumoniae* BNCC 353393 bacteria suspensions in 0.9% saline. This provides a challenge inoculum of approximately $1 \times 10^6$ cells per thigh. At 2 hours post infection, mice were given 100 μL of each antibiotic at different concentrations via SC

injection. At 24 hours post infection, the mice were euthanized and their thigh muscles were aseptically removed, weighed, homogenized, and enumerated for bacterial burden by colony forming unit (CFU) counts. All graphical data were presented as data points (n=3 mice/n= 6 thighs) by group and were statistically analyzed using Prism 9.0. All animal study procedures were approved by the Animal Ethics Committee of China Pharmaceutical University (Approved number: 2024-08-078).

**Neutropenic skin infection model:**

BALB/c mice weighing 20-22 g were used in this study, half male and half female (Hangzhou Medical College, China). The mice were randomly grouped in individual cages and acclimatized for three days before the experiments. To induce neutropenia, the mice were injected with 50 mg/kg of cyclophosphamide via intraperitoneal injection on the third and first days before infection(*7, 8*). Subsequently, mice were anesthetized with 4% chloral hydrate (80 μL/10 g body weight) via intraperitoneal injection. Then the hair on the mice's backs were removed using a razor, and full-thickness skin perforations were made on the dorsal skin using a biopsy puncher with a diameter of 1.0 cm. A suspension of *S. aureus* ATCC BAA-44 at a concentration of $2.4\times10^6$ CFU/mL was evenly dripped on the wound (20 μL/ mice). The mounds were topically treated with vancomycin (20 mg/kg) and paenimycin (1 mg/kg, 10 mg/kg, 20 mg/kg and 40 mg/kg), respectively. All compounds were administered via subcutaneous injection once daily for five consecutive days. Photographic records were taken on days 1, 3, 5, 8, 10, 12 and 14 post infection. On day 15, the wound tissues were collected and the bacterial burden was recorded using the plate counting method. All graphical data were presented as data points (n=5 mice) by group and were statistically analyzed using Prism 9.0. All animal study procedures were approved by the Animal Ethics Committee of China Pharmaceutical University (Approved number. 2024-10-112).

**Neutropenic vaginal infection model:**

Female BALB/c mice weighing 22-25 g were used in this study (Hangzhou Medical College, China). The mice were randomly grouped (n=5, each group) and acclimatized for three days before the experiment. The mice were subcutaneously injected with 17-β-estradiol (25 mg/kg) for three days before *Neisseria gonorrhoeae bacteria* challenge(*9-11*). On the first day of 17-β-estradiol administration, mice were injected with cyclophosphamide (50 mg/kg, i.p.). For the subsequent two days, antibiotics (Vancomycin+Streptomycin) were administered intraperitoneally once daily to suppress the overgrowth of commensal bacteria in the murine reproductive tract. On the day of infection, physical damage to the vaginal mucosa was induced using a cotton swab. Then, 50 μL of *Neisseria gonorrhoeae* ATCC 31426 suspension ($5\times10^8$ CFU/mL) was inoculated into the vaginal cavity using a pipette and subjected tail suspension for 5 min. This infection procedure was repeated for three consecutive days.

On the second day after the last infection, mice were subcutaneously injected with paenimycin at varying doses (1 mg/kg, 5 mg/kg, 10 mg/kg and 20 mg/kg) once daily for three consecutive days. On the second day after the last administration, mouse vaginal secretions were smeared and examined under a microscope. Additionally, vaginal lavage fluid were collected (about 200 μL), and the bacterial burden was recorded. All graphical data were presented as data points by group and were statistically analyzed using Prism 9.0. All animal study procedures were approved by the Animal Ethics Committee of China Pharmaceutical University (Approved number. 2024-10-112).

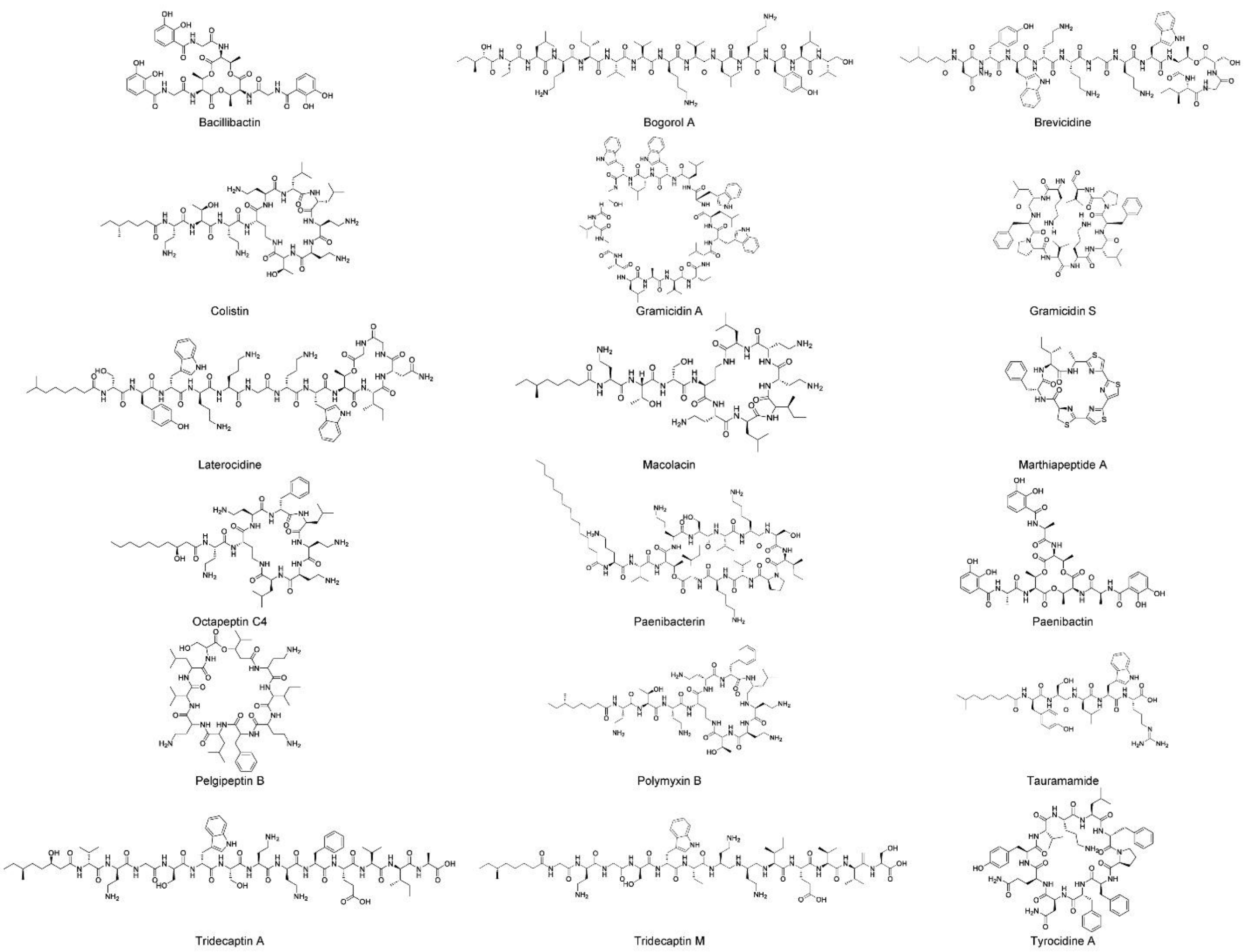


**Fig. S1:** Structures of some nonribosomal peptides (NRPs) natural products from *Paenibacillaceae* family.

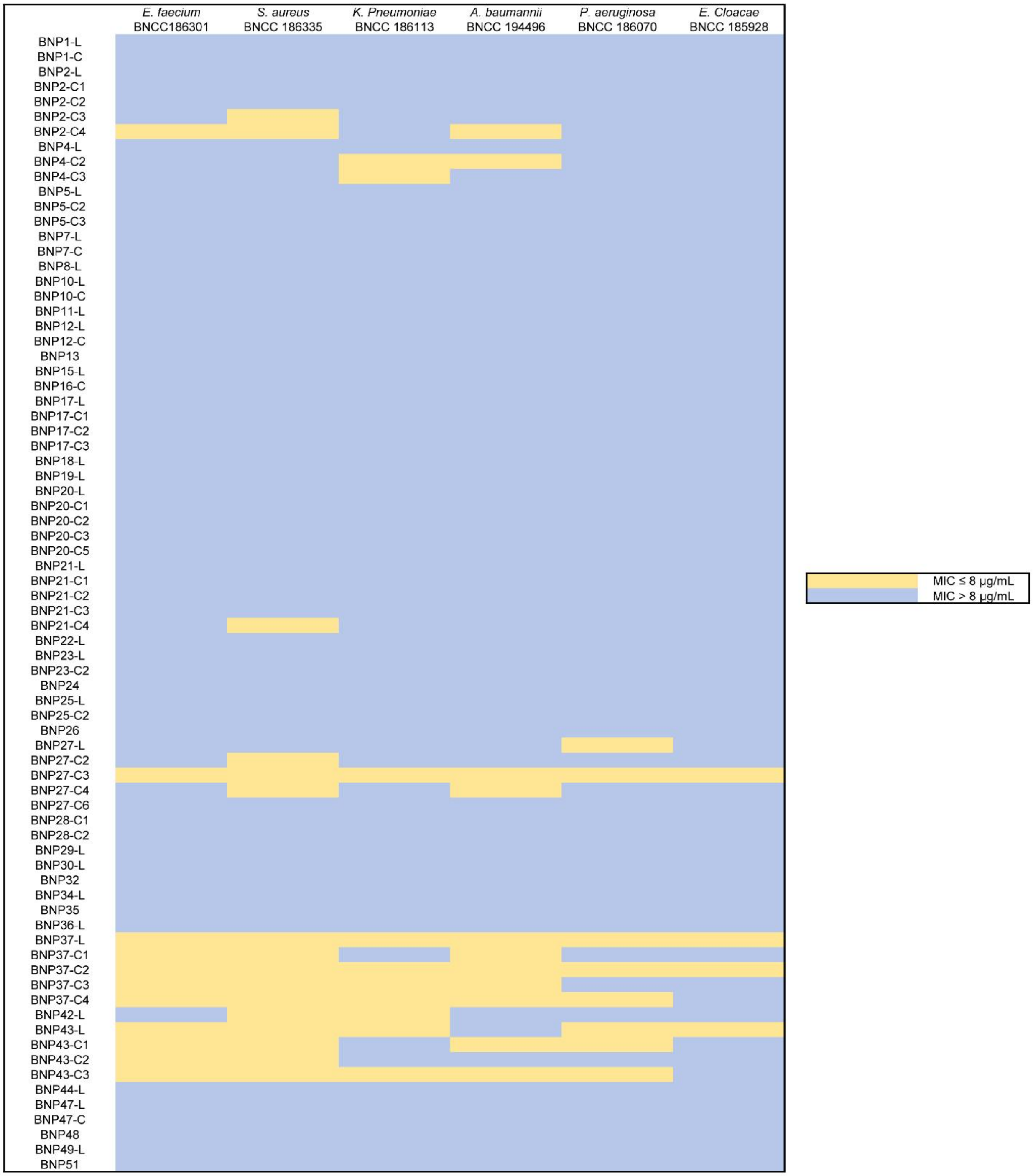


**Fig. S2:** Bioactivity screening of all synthesized peptides. MIC values ≤ 8 μg/mL were marked in yellow and MIC values > 8 μg/mL were marked in blue.

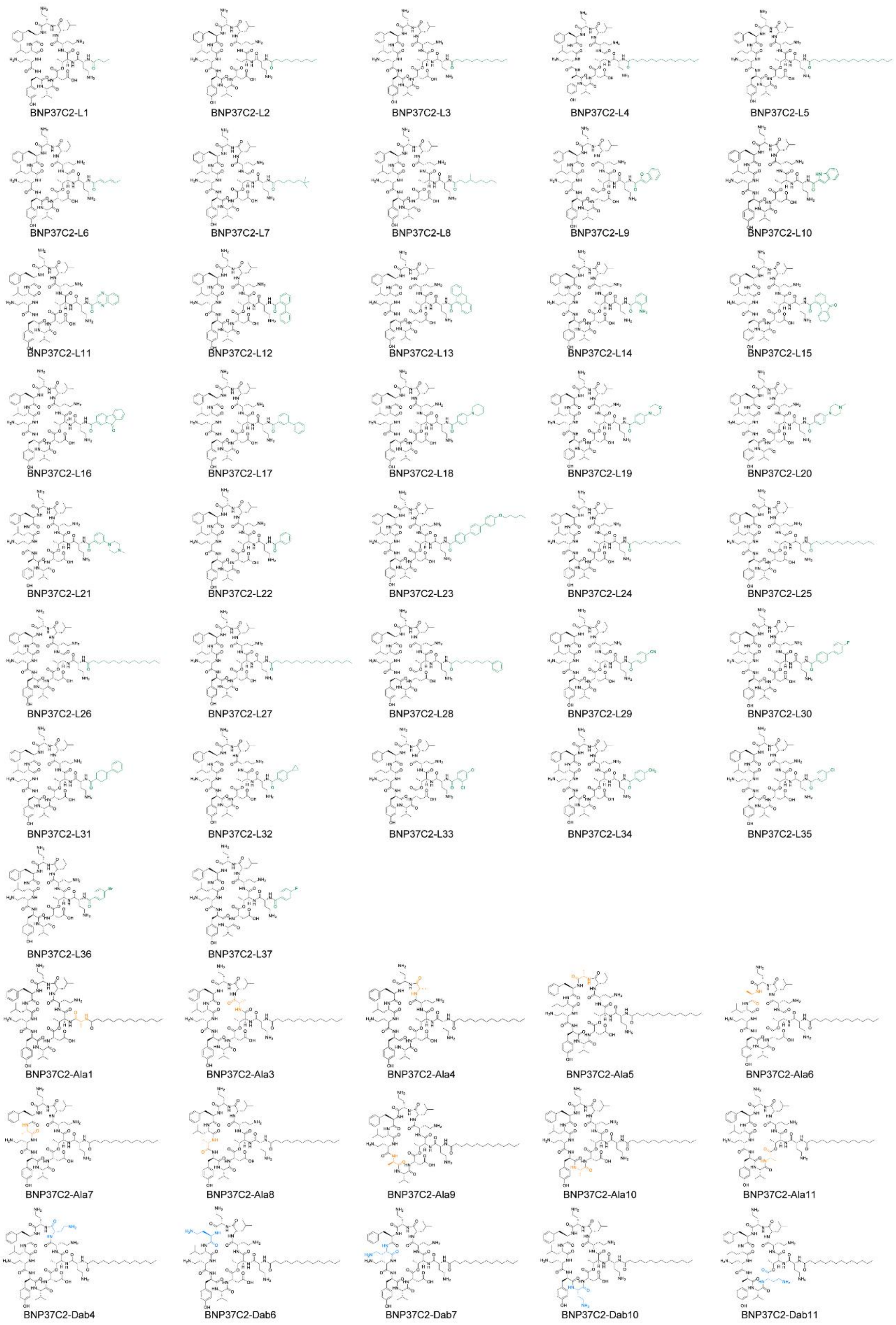


**Fig. S3:** Structures of paenimycin analogs.

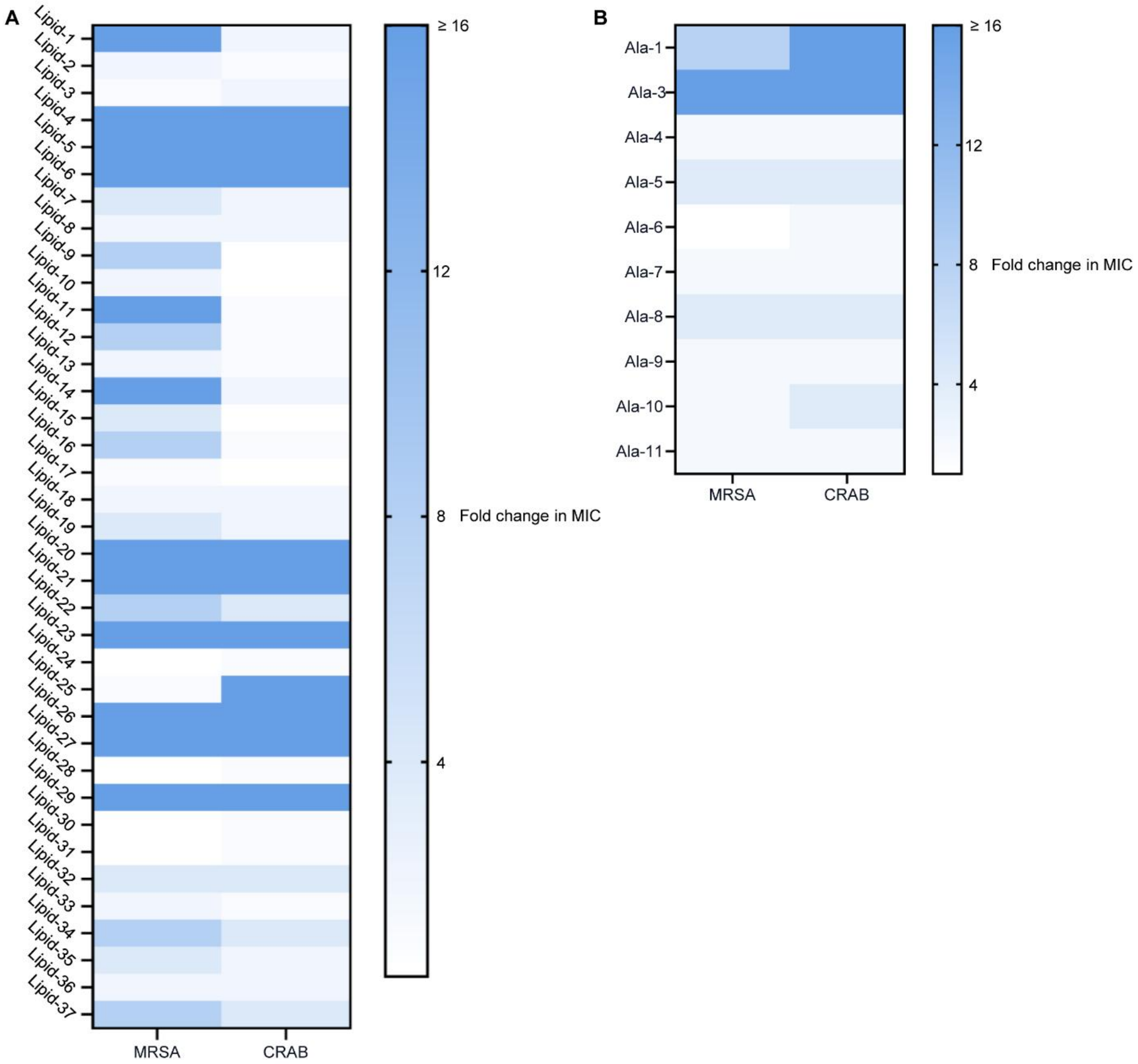


**Fig. S4:** Structure optimization of BNP37C2. A) Fold changes in MIC after lipid screening. B) Fold changes in MIC after Ala screening. Note: MRSA (methicillin-resistant *S. aureus*, *S. aureus* ATCC BAA44) and CRAB (carbapenems-resistant *A. baumannii*, *A. baumannii* ATCC BAA1605).

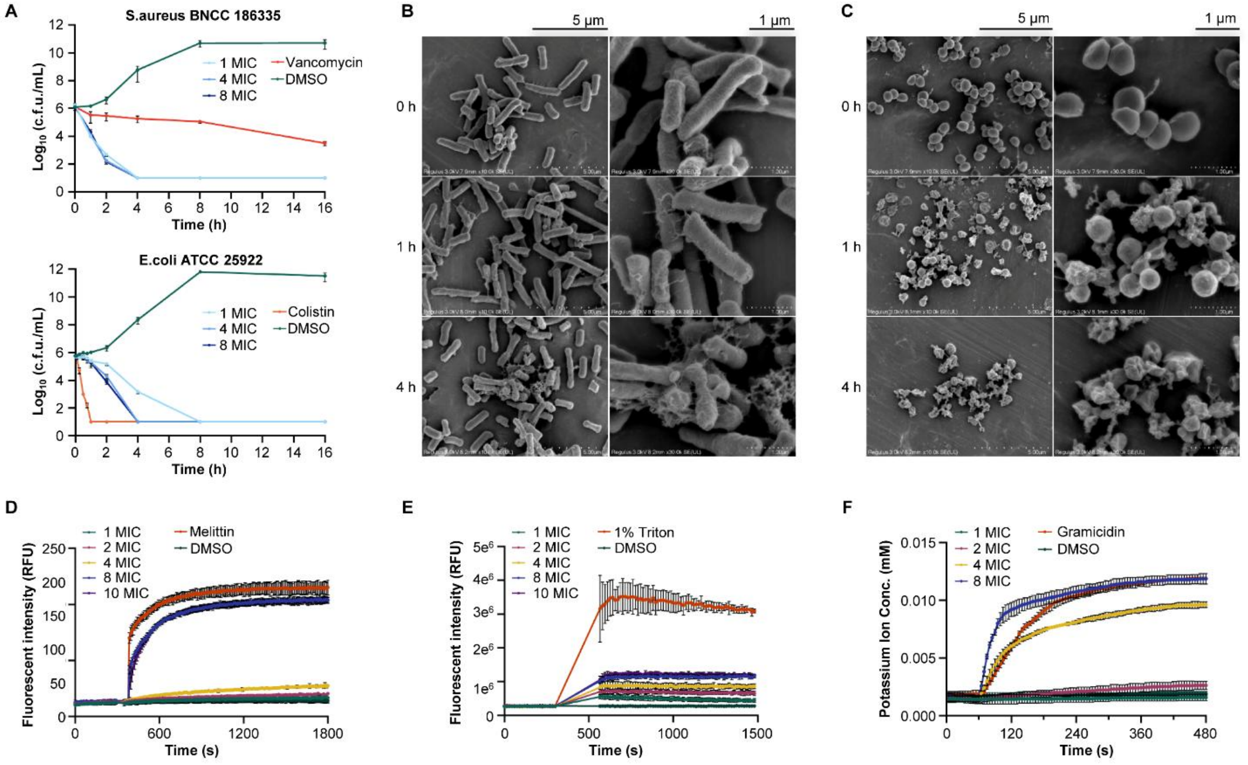


**Fig. S5:** Paenimycin exhibits bactericidal activity. A) Time-dependent killing curve of paenimycin against *S. aureus* BNCC 186335 and *E. coli* ATCC 25922. Bacteria were treated with 1, 4, 8×MIC of paenimycin and 8× MIC of vancomycin or colistin, respectively. CFUs (colony-forming units) were counted three independent times. B) Scanning electron microscope (SEM) image of *E. coli* ATCC 25922 treated with 8× MIC of paenimycin for 0,1 and 4 hours. C) SEM image of *S. aureus* BNCCC 186335 treated with 8×MIC of paenimycin for 0,1 and 4 hours. D) Cell lysis was tested by SYTOX dye. *S. aureus* BNCC 186335 was treated with 1, 2, 4, 8, 10×MIC of paenimycin and 10×MIC of melittin was used as a lysis control. E) Membrane depolarization was detected by DiSC3(5) dye. DMSO was used as a depolarized control, and *S. aureus* BNCC 186335 was treated with 1, 2, 4, 8, 10×MIC of paenimycin and 1% Triton X. F) Potassium release effect of paenimycin. 8×MIC of gramicidin was used as a control, and the concentration of potassium ions were measured after treating with 1, 2, 4, 8×MIC of paenimycin

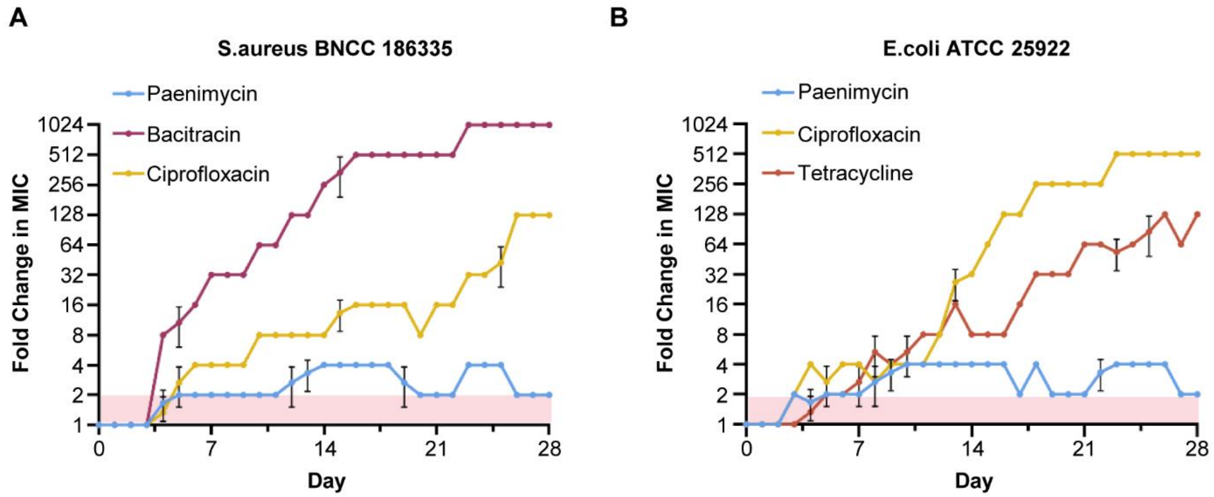


**Fig. S6:** Paenimycin showed no resistance under lab conditions. A) Daily serial passaging of *S. aureus* BNCC 186335 at sub-MIC concentrations of paenimycin, bacitracin and ciprofloxacin. B) *E. coli* ATCC 25922 at sub-MIC concentrations of paenimycin, ciprofloxacin and tetracycline.

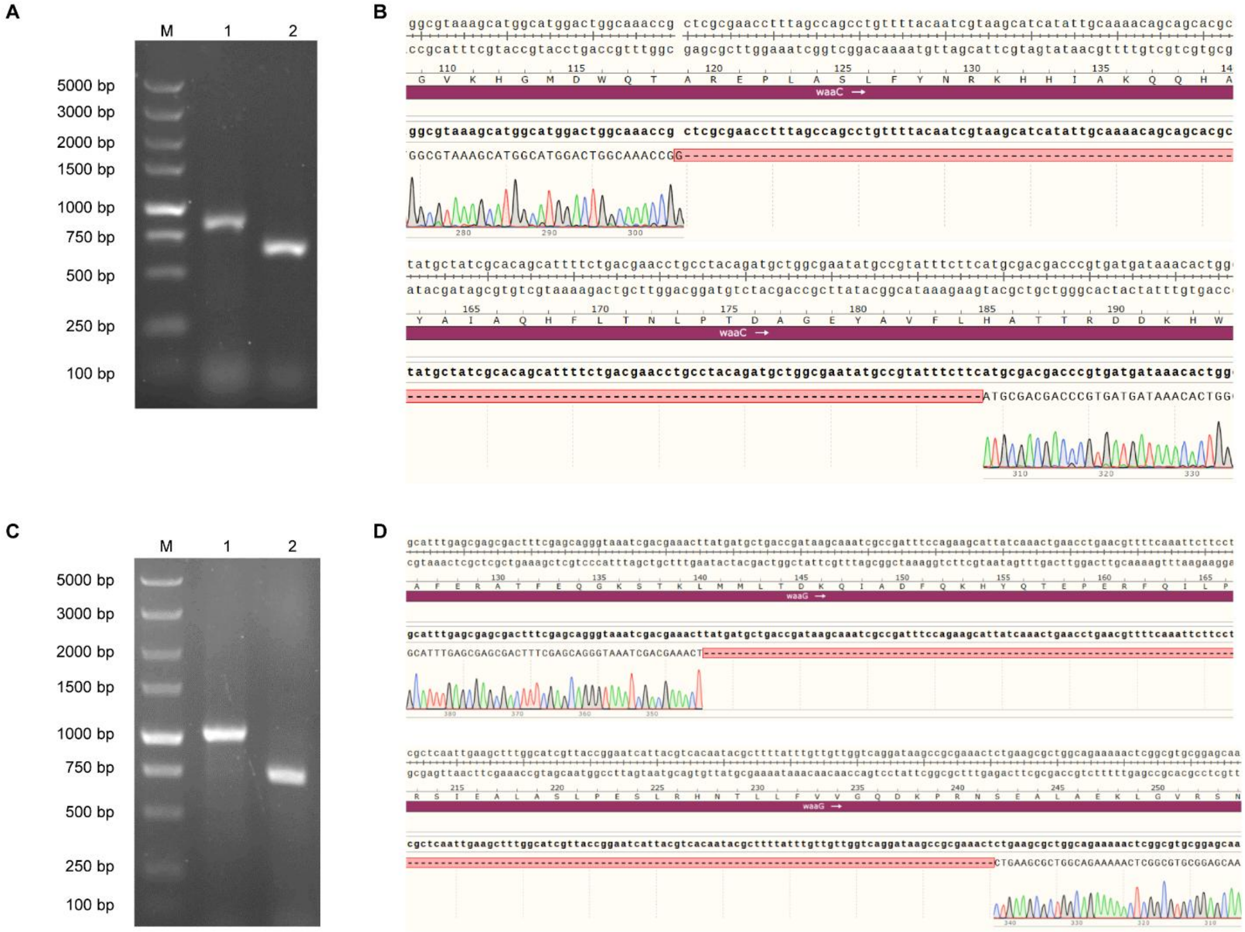


**Fig. S7:** A) PCR amplification of the knockout region in *waaC*. Lane 1: wide type (WT) Lane 2: Δ *waaC*; M, *GL* DNA Marker 5000. B) Sequencing of the Δ *waaC* mutant confirms the deletion of 197 bp region. C) PCR amplification of the knockout region in *waaG*. Lane 1: wide type (WT) Lane 2: Δ *waaG* ; M, *GL* DNA Marker 5000. D) Sequencing of the Δ *waaG* mutant confirms the deletion of 305 bp region.

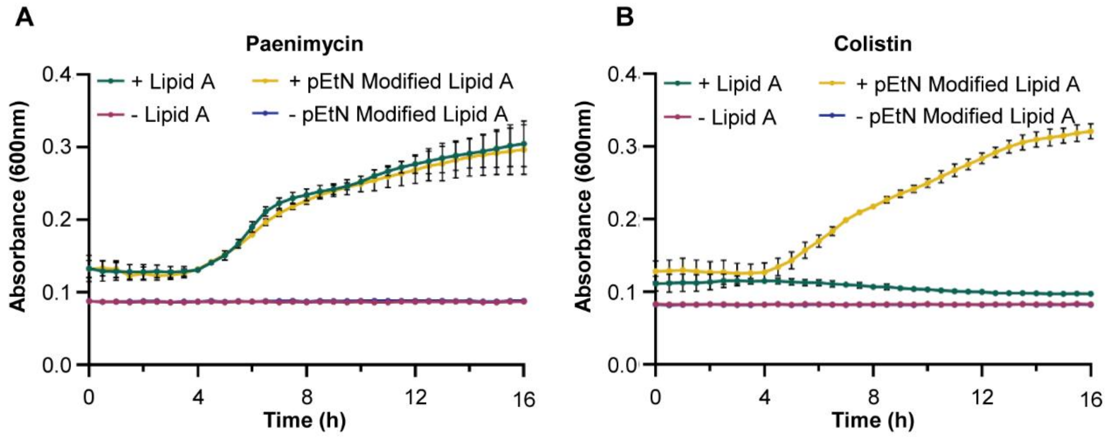


**Fig. S8:** Inhibition of *E. coli* ATCC 25922 growths by paenimycin and colistin after addition of extracted pEtN modified and unmodified lipid A. Growth medium-insoluble lipid A leads to increased initial absorbance values.

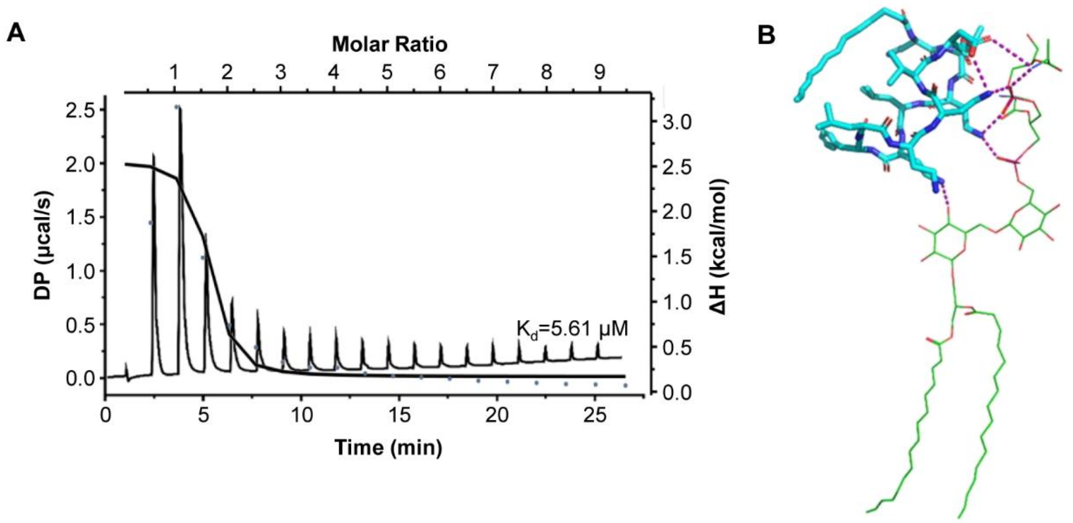


**Fig. S9:** Paenimycin and colistin binding to LPS or LTA. A) ITC assay result of paenimycin binding with LTA. B) MD simulation results of paenimycin with LTA.

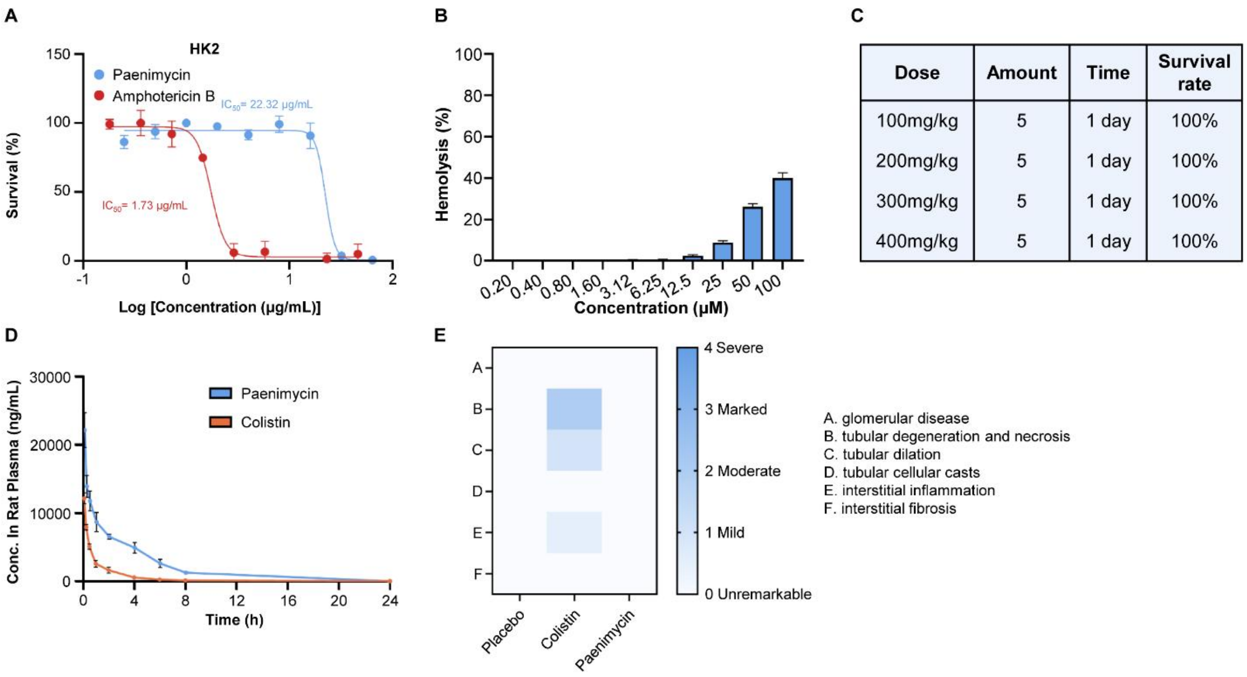


| Dose | Amount | Time | Survival rate |
|---|---|---|---|
| 100mg/kg | 5 | 1 day | 100% |
| 200mg/kg | 5 | 1 day | 100% |
| 300mg/kg | 5 | 1 day | 100% |
| 400mg/kg | 5 | 1 day | 100% |

**Fig. S10:** *In vivo* and *in vitro* safety assays of paenimycin. A) Cytotoxicity pf paenimycin against HK2 cell line. Amphotericin B was used as a control. B) Hemolysis assay of paenimycin. C) Most tolerant assay of paenimycin (s.c. qd, n=5 per group). D) The pharmacokinetic curves of paenimycin and colistin over time after administration of 5 mg/kg via intravenous injection. E) Histopathological analysis of kidney sections collected 7 days after mice treated with vehicle, colistin and paenimycin. (s.c. qd, n=3 per group).

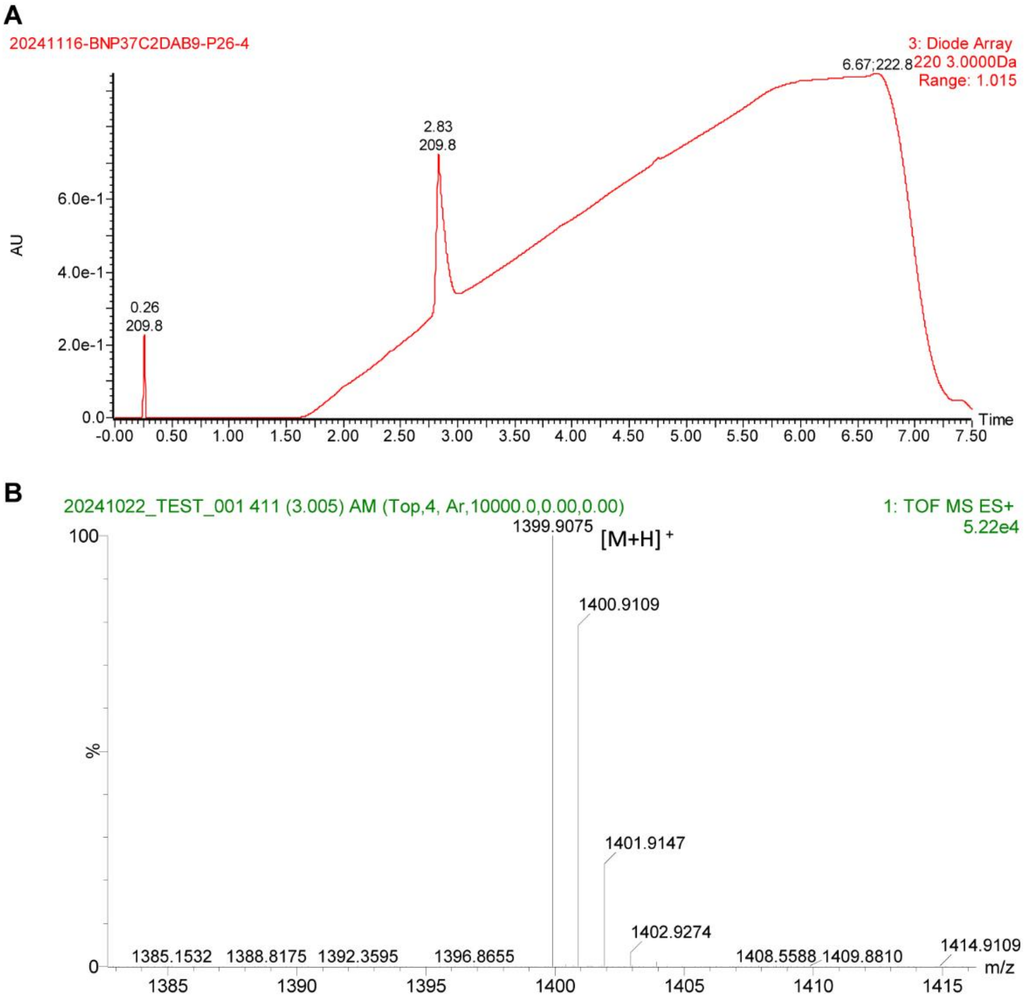


**Fig. S11:** LCMS analysis of paenimycin. A) HPLC chromatogram of paenimycin. B) High-resolution mass spectra (HRMS) of paenimycin.

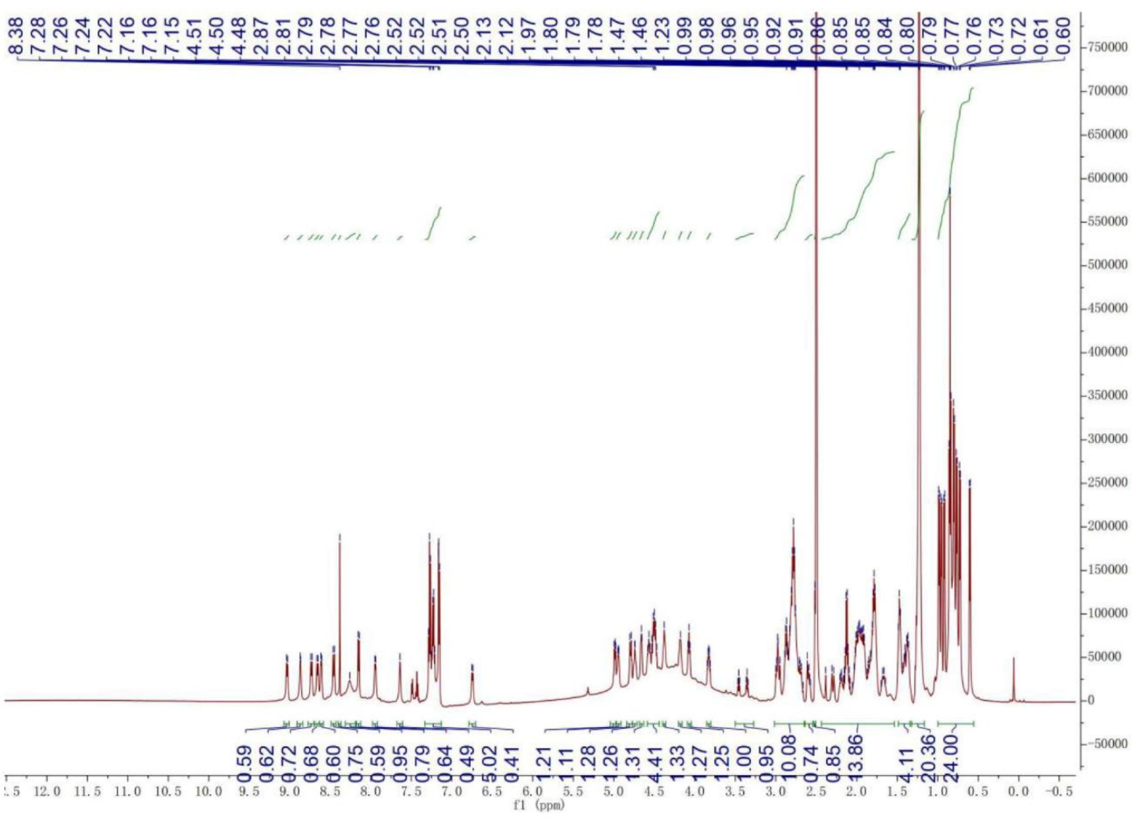

**Fig. S12:** $^{1}$H NMR (*d6*-DMSO, 600 MHz) spectrum of paenimycin.

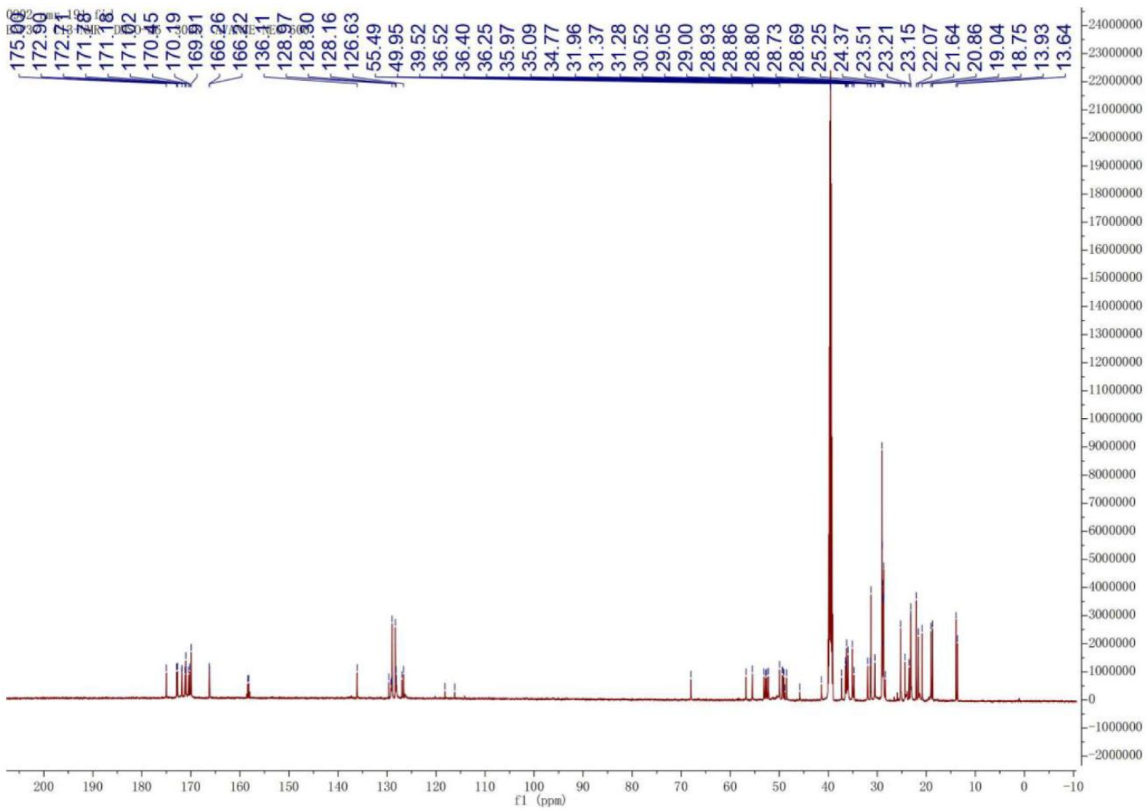

**Fig. S13:** $^{13}$C NMR (*d6*-DMSO, 600 MHz) spectrum of paenimycin.

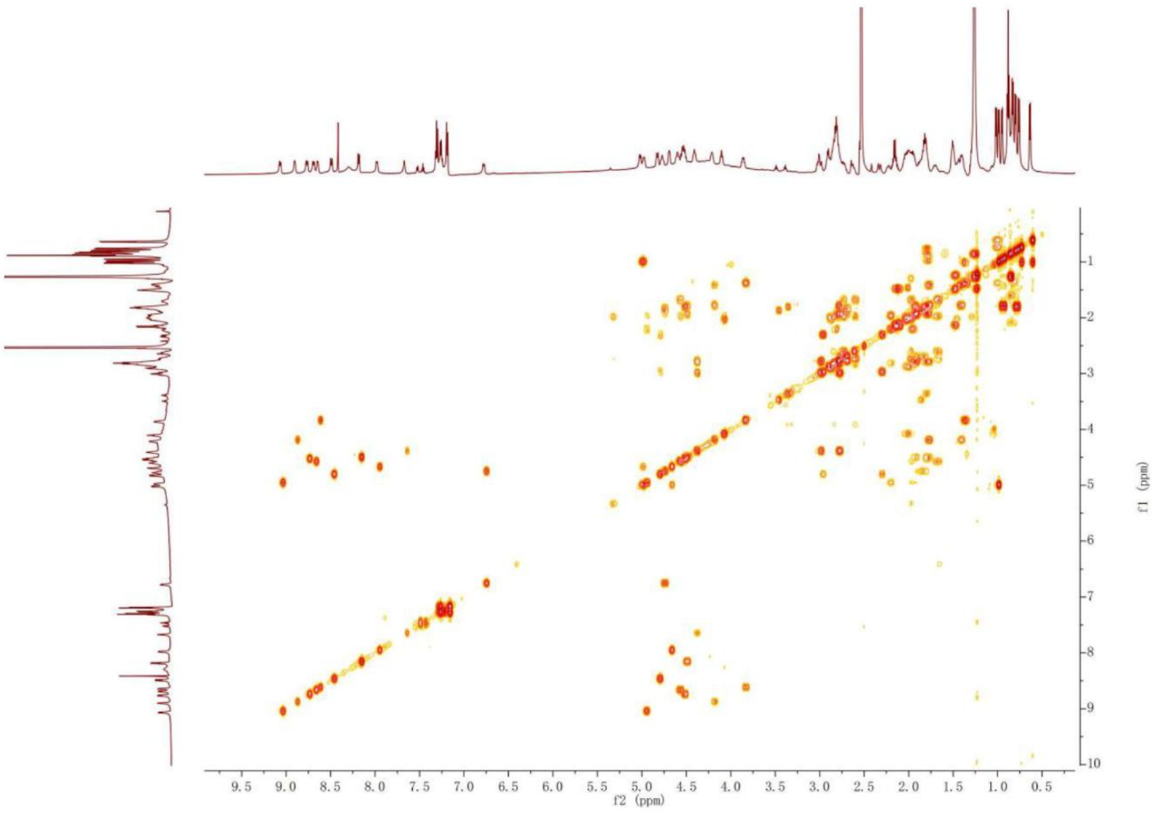


**Fig. S14:** COSY NMR (*d6*-DMSO, 600 MHz) spectrum of paenimycin.

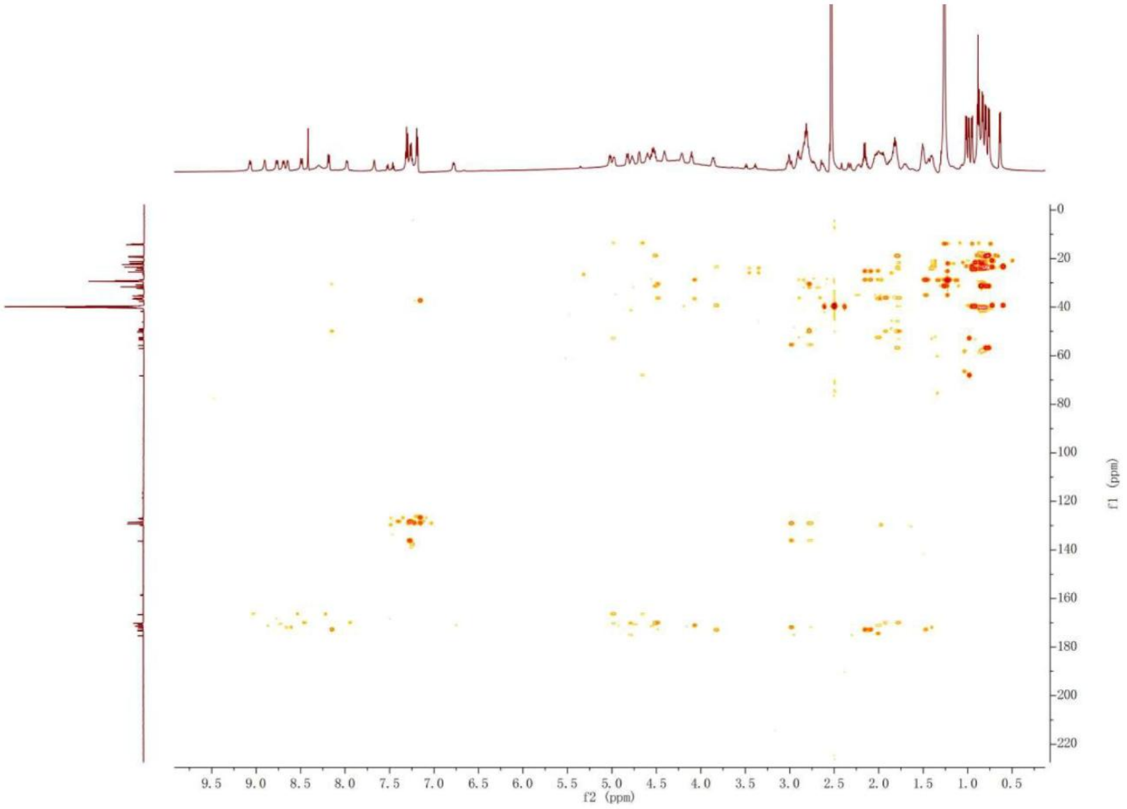


**Fig. S15:** HMBC NMR (*d6*-DMSO, 600 MHz) spectrum of paenimycin.

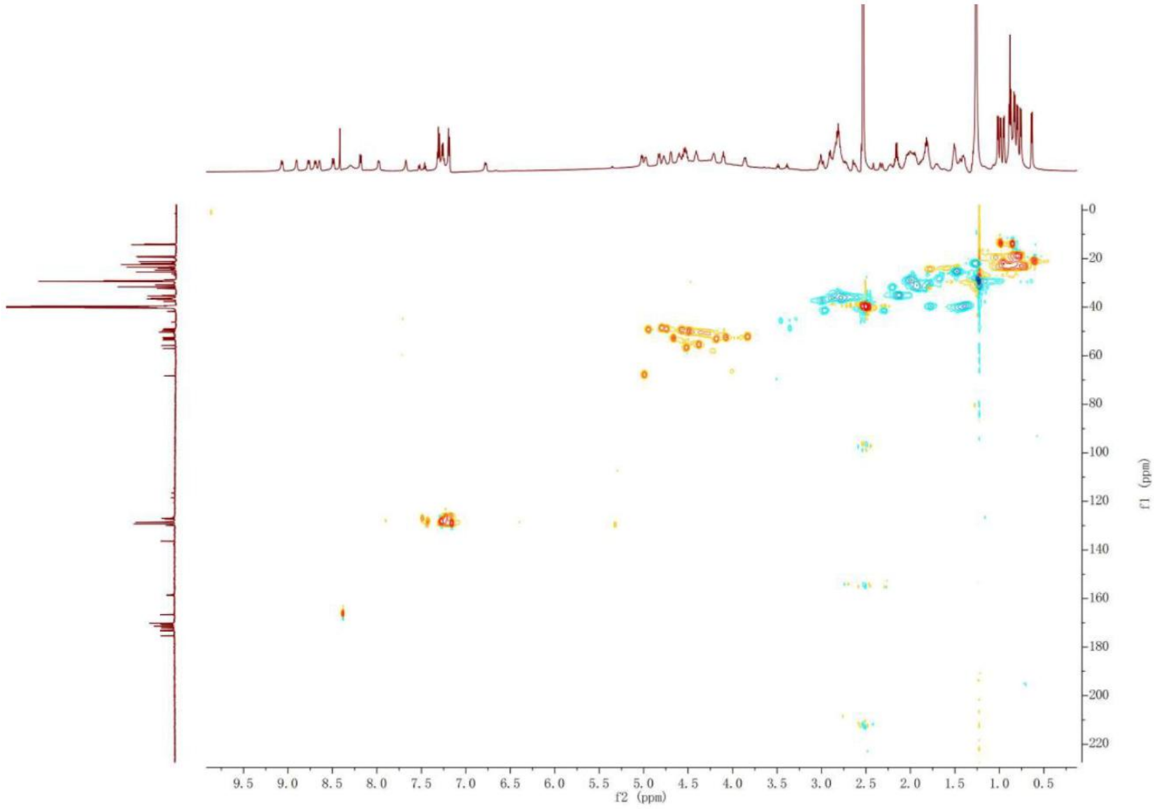


**Fig. S16:** HSQC NMR (*d6*-DMSO, 600 MHz) spectrum of paenimycin.

**Table S1:** *Pae* biosynthetic gene cluster annotations (GenBank Accession Number: NZ_JAQAGY010000015.1 location: 1-74,669).

| ORF | Gene name | Gene size (bp) | Proposed function | Protein, [Source Organism}, Accession number |
|---|---|---|---|---|
| 1 | *paeA* | 1287 | Regulator | FAD-dependent oxidoreductase, [Paenibacillus caseinilyticus], WP_269868027.1 |
| 2 | *paeB* | 12306 | Biosynthesis gene | Non-ribosomal peptide synthetase, [Paenibacillus caseinilyticus] |
| 3 | *paeC* | 25509 | Biosynthesis gene | Non-ribosomal peptide synthetase, [Paenibacillus caseinilyticus] |
| 4 | *paeD* | 3279 | Biosynthesis gene | Non-ribosomal peptide synthetase, [Paenibacillus caseinilyticus] |
| 5 | *paeE* | 1719 | Transpoter | ABC transporter ATP-binding protein, [Paenibacillus caseinilyticus], WP_269868034.1 |
| 6 | *paeF* | 1797 | Transporter | ABC transporter ATP-binding protein, [Paenibacillus caseinilyticus], WP_269868033.1 |

**Table S2:** High-resolution mass spectrometry data for candidate peptides.

| Molecules name | Chemical Formula | Theoretical Mass | Observed mass | Error (ppm) |
|---|---|---|---|---|
| **BNP1-L** | $C_{69}H_{114}N_{14}O_{16}$ | $[M+H]^+$ 1395.8615 | $[M+H]^+$ 1395.8550 | -4.65 |
| **BNP1-C** | $C_{69}H_{112}N_{14}O_{15}$ | $[M+H]^+$ 1377.8510 | $[M+H]^+$ 1377.8475 | -2.54 |
| **BNP2-L** | $C_{100}H_{138}N_{16}O_{24}$ | $[M+H]^+$ 1948.0148 | $[M+H]^+$ 1948.0211 | 3.23 |
| **BNP2-C1** | $C_{100}H_{136}N_{16}O_{23}$ | $[M+H]^+$ 1930.0042 | $[M+H]^+$ 1930.0016 | -1.38 |
| **BNP2-C2** | $C_{100}H_{136}N_{16}O_{23}$ | $[M+H]^+$ 1930.0042 | $[M+H]^+$ 1930.0111 | 3.58 |
| **BNP2-C3** | $C_{100}H_{136}N_{16}O_{23}$ | $[M+H]^+$ 1930.0042 | $[M+H]^+$ 1930.0016 | -1.38 |
| **BNP2-C4** | $C_{100}H_{136}N_{16}O_{23}$ | $[M+H]^+$ 1930.0042 | $[M+H]^+$ 1930.0016 | -1.38 |
| **BNP4-L** | $C_{58}H_{96}N_{14}O_{16}$ | $[M+H]^+$ 1245.7207 | $[M+H]^+$ 1245.7238 | 2.49 |
| **BNP4-C2** | $C_{58}H_{94}N_{14}O_{15}$ | $[M+H]^+$ 1227.7101 | $[M+H]^+$ 1227.7164 | 5.13 |
| **BNP4-C3** | $C_{58}H_{94}N_{14}O_{15}$ | $[M+H]^+$ 1227.7101 | $[M+H]^+$ 1227.7144 | 3.50 |
| **BNP5-L** | $C_{79}H_{128}N_{16}O_{17}$ | $[M+H]^+$ 1573.9722 | $[M+H]^+$ 1573.9800 | 4.96 |
| **BNP5-C2** | $C_{79}H_{126}N_{16}O_{16}$ | $[M+H]^+$ 1555.9616 | $[M+H]^+$ 1555.9597 | 1.20 |
| **BNP5-C3** | $C_{79}H_{126}N_{16}O_{16}$ | $[M+H]^+$ 1555.9616 | $[M+H]^+$ 1555.9655 | 2.51 |
| **BNP7-L** | $C_{40}H_{68}N_8O_9$ | $[M+H]^+$ 805.5188 | $[M+H]^+$ 805.5188 | 0.00 |
| **BNP7-C** | $C_{40}H_{66}N_8O_8$ | $[M+H]^+$ 787.5082 | $[M+H]^+$ 787.5022 | -7.61 |
| **BNP8-L** | $C_{63}H_{95}N_7O_{13}$ | $[M+H]^+$ 1158.7066 | $[M+H]^+$ 1158.7098 | 2.76 |
| **BNP10-L** | $C_{50}H_{88}N_{10}O_9S$ | $[M+H]^+$ 1005.6535 | $[M+H]^+$ 1005.6464 | -7.06 |
| **BNP10-C** | $C_{50}H_{86}N_{10}O_8S$ | $[M+H]^+$ 987.6429 | $[M+H]^+$ 987.6407 | -2.23 |
| **BNP11-L** | $C_{65}H_{108}N_{10}O_{15}$ | $[M+H]^+$ 1269.8074 | $[M+H]^+$ 1269.8109 | 2.75 |
| **BNP12-L** | $C_{50}H_{71}N_9O_{11}$ | $[M+H]^+$ 974.5351 | $[M+H]^+$ 974.5355 | 0.41 |
| **BNP12-C** | $C_{50}H_{69}N_9O_{10}$ | $[M+H]^+$ 956.5246 | $[M+H]^+$ 956.5187 | -6.12 |
| **BNP13** | $C_{57}H_{90}N_{10}O_{12}$ | $[M+H]^+$ 1107.6818 | $[M+H]^+$ 1107.6838 | 1.81 |
| **BNP15-L** | $C_{61}H_{86}N_{12}O_{14}$ | $[M+H]^+$ 1211.6465 | $[M+H]^+$ 1211.6547 | 6.77 |
| **BNP16-C** | $C_{36}H_{63}N_7O_8$ | $[M+H]^+$ 722.4818 | $[M+H]^+$ 722.4818 | 0.00 |
| **BNP17-L** | $C_{46}H_{75}N_9O_{14}$ | $[M+H]^+$ 978.5512 | $[M+H]^+$ 978.5522 | 1.02 |
| **BNP17-C1** | $C_{46}H_{73}N_9O_{13}$ | $[M+H]^+$ 960.5406 | $[M+H]^+$ 960.5433 | 2.81 |
| **BNP17-C2** | $C_{46}H_{73}N_9O_{13}$ | $[M+H]^+$ 960.5406 | $[M+H]^+$ 960.5419 | 1.35 |
| **BNP17-C3** | $C_{46}H_{73}N_9O_{13}$ | $[M+H]^+$ 960.5406 | $[M+H]^+$ 960.5427 | 2.19 |
| **BNP18-L** | $C_{56}H_{88}N_{10}O_{12}$ | $[M+H]^+$ 1093.6661 | $[M+H]^+$ 1093.6678 | 1.55 |
| **BNP19-L** | $C_{41}H_{63}N_7O_7$ | $[M+H]^+$ 766.4867 | $[M+H]^+$ 766.4871 | 0.52 |
| **BNP20-L** | $C_{54}H_{90}N_{12}O_{17}$ | $[M+H]^+$ 1179.6625 | $[M+H]^+$ 1179.6556 | -5.84 |
| **BNP20-C1** | $C_{54}H_{88}N_{12}O_{16}$ | $[M+H]^+$ 1161.6519 | $[M+H]^+$ 1161.6418 | -8.69 |
| **BNP20-C2** | $C_{54}H_{88}N_{12}O_{16}$ | $[M+H]^+$ 1161.6519 | $[M+H]^+$ 1161.6542 | 1.98 |

| | | | | |
|---|---|---|---|---|
| **BNP20-C3** | C54H88N12O16 | $[M+H]^+$ 1161.6519 | $[M+H]^+$ 1161.6530 | 0.95 |
| **BNP20-C5** | C54H88N12O16 | $[M+H]^+$ 1161.6519 | $[M+H]^+$ 1161.6543 | 2.07 |
| **BNP21-L** | C90H135N15O22 | $[M+H]^+$ 1778.9984 | $[M+H]^+$ 1779.0039 | 3.09 |
| **BNP21-C1** | C90H133N15O21 | $[M+H]^+$ 1760.9879 | $[M+H]^+$ 1760.9938 | 3.35 |
| **BNP21-C2** | C90H133N15O21 | $[M+H]^+$ 1760.9879 | $[M+H]^+$ 1760.9923 | 2.50 |
| **BNP21-C3** | C90H133N15O21 | $[M+H]^+$ 1760.9879 | $[M+H]^+$ 1760.9938 | 3.35 |
| **BNP21-C4** | C90H133N15O21 | $[M+H]^+$ 1760.9879 | $[M+H]^+$ 1760.9734 | -8.23 |
| **BNP22-L** | C52H89N9O12 | $[M+H]^+$ 1032.6709 | $[M+H]^+$ 1032.6730 | 2.00 |
| **BNP23-L** | C78H124N16O22 | $[M+2H]^{2+}$ 819.4616 | $[M+2H]^{2+}$ 819.4630 | 1.70 |
| **BNP23-C2** | C78H122N16O21 | $[M+2H]^{2+}$ 810.4563 | $[M+2H]^{2+}$ 810.4509 | -6.66 |
| **BNP24** | C40H69N9O14 | $[M+H]^+$ 900.5042 | $[M+H]^+$ 900.5023 | -2.11 |
| **BNP25-L** | C46H77N7O14 | $[M+H]^+$ 952.5607 | $[M+H]^+$ 952.5629 | 2.31 |
| **BNP25-C2** | C46H75N7O13 | $[M+H]^+$ 934.5501 | $[M+H]^+$ 934.5511 | 1.07 |
| **BNP26** | C45H83N7O10 | $[M+H]^+$ 882.6280 | $[M+H]^+$ 882.6304 | 2.72 |
| **BNP27-L** | C87H155N19O20 | $[M+2H]^{2+}$ 894.0926 | $[M+2H]^{2+}$ 894.0904 | 1.71 |
| **BNP27-C2** | C87H153N19O19 | $[M+H]^+$ 1769.1668 | $[M+H]^+$ 1769.1644 | 1.47 |
| **BNP27-C3** | C87H153N19O19 | $[M+H]^+$ 1769.1668 | $[M+H]^+$ 1769.1644 | 1.47 |
| **BNP27-C4** | C87H153N19O19 | $[M+H]^+$ 1769.1668 | $[M+H]^+$ 1769.1740 | 4.07 |
| **BNP27-C6** | C87H153N19O19 | $[M+H]^+$ 1769.1668 | $[M+H]^+$ 1769.1644 | 1.47 |
| **BNP28-C1** | C72H105N15O17 | $[M+H]^+$ 1452.7891 | $[M+H]^+$ 1452.7781 | -7.57 |
| **BNP28-C2** | C72H105N15O17 | $[M+H]^+$ 1452.7891 | $[M+H]^+$ 1452.7781 | -7.57 |
| **BNP29-L** | C49H75N7O15 | $[M+H]^+$ 1002.5399 | $[M+H]^+$ 1002.5286 | -1.13 |
| **BNP30-L** | C60H90N8O10 | $[M+H]^+$ 1083.6858 | $[M+H]^+$ 1083.6746 | -1.03 |
| **BNP32** | C45H81N7O12 | $[M+H]^+$ 912.6021 | $[M+H]^+$ 912.6069 | 5.26 |
| **BNP34-L** | C42H73N9013 | $[M+H]^+$ 912.5406 | $[M+H]^+$ 912.5366 | -4.38 |
| **BNP35** | C67H98N8O13 | $[M+H]^+$ 1223.7332 | $[M+H]^+$ 1223.7368 | 2.94 |
| **BNP36-L** | C43H77N7O13 | $[M+H]^+$ 900.5658 | $[M+H]^+$ 900.5648 | -1.11 |
| **BNP37-L** | C73H121N15O17 | $[M+H]^+$ 1480.9143 | $[M+H]^+$ 1480.9229 | 5.82 |
| **BNP37-C1** | C73H119N15O16 | $[M+H]^+$ 1462.9037 | $[M+H]^+$ 1462.9012 | -1.71 |
| **BNP37-C2** | C73H119N15O16 | $[M+H]^+$ 1462.9037 | $[M+H]^+$ 1462.9117 | 5.47 |
| **BNP37-C3** | C73H119N15O16 | $[M+H]^+$ 1462.9037 | $[M+H]^+$ 1462.9117 | 5.47 |
| **BNP37-C4** | C73H119N15O16 | $[M+H]^+$ 1462.9037 | $[M+H]^+$ 1462.9020 | -1.16 |
| **BNP42-L** | C54H91N9O9 | $[M+H]^+$ 1010.7018 | $[M+H]^+$ 1010.6969 | -4.85 |
| **BNP43-L** | C56H106N12O14 | $[M+H]^+$ 1171.8030 | $[M+H]^+$ 1171.7939 | -7.77 |
| **BNP43-C1** | C56H104N12O13 | $[M+H]^+$ 1153.7924 | $[M+H]^+$ 1153.7865 | -5.11 |

| | | | | |
|---|---|---|---|---|
| **BNP43-C2** | $C_{56}H_{104}N_{12}O_{13}$ | $[M+H]^+$ 1153.7924 | $[M+H]^+$ 1153.7935 | 0.95 |
| **BNP43-C3** | $C_{56}H_{104}N_{12}O_{13}$ | $[M+H]^+$ 1153.7924 | $[M+H]^+$ 1153.7935 | 0.95 |
| **BNP44-L** | $C_{45}H_{75}N_7O_{14}$ | $[M+H]^+$ 938.5450 | $[M+H]^+$ 938.5457 | 0.75 |
| **BNP47-L** | $C_{63}H_{97}N_9O_9$ | $[M+H]^+$ 1124.7488 | $[M+H]^+$ 1124.7548 | 5.33 |
| **BNP47-C** | $C_{63}H_{95}N_9O_8$ | $[M+H]^+$ 1106.7382 | $[M+H]^+$ 1106.7424 | 3.80 |
| **BNP48** | $C_{51}H_{93}N_9O_{12}$ | $[M+H]^+$ 1024.7022 | $[M+H]^+$ 1024.6947 | -7.32 |
| **BNP49-L** | $C_{36}H_{63}N_5O_{11}$ | $[M+H]^+$ 742.4602 | $[M+H]^+$ 742.4632 | 4.04 |
| **BNP51** | $C_{34}H_{59}N_9O_{13}$ | $[M+H]^+$ 802.4311 | $[M+H]^+$ 802.4327 | 1.99 |

**Table S3:** High-resolution mass spectrometry data for paenimycin analogs.

| Molecules name | Chemical Formula | Theoretical Mass | Observed mass | Error (ppm) |
|---|---|---|---|---|
| **BNP37C2-L1** | C63H99N15O16 | $[M+H]^+$ 1322.7472 | $[M+H]^+$ 1322.7446 | 1.97 |
| **BNP37C2-L2** | C69H111N15O16 | $[M+H]^+$ 1406.8411 | $[M+H]^+$ 1406.8414 | 0.21 |
| **BNP37C2-L3** | C71H115N15O16 | $[M+H]^+$ 1434.8724 | $[M+H]^+$ 1434.8774 | 3.48 |
| **BNP37C2-L4** | C75H123N15O16 | $[M+H]^+$ 1490.9350 | $[M+H]^+$ 1490.9395 | 3.02 |
| **BNP37C2-L5** | C77H127N15O16 | $[M+H]^+$ 1518.9633 | $[M+H]^+$ 1518.9666 | 2.17 |
| **BNP37C2-L6** | C65H99N15O16 | $[M+2H]^{2+}$ 673.8775 | $[M+2H]^{2+}$ 673.8749 | -3.86 |
| **BNP37C2-L7** | C69H111N15O16 | $[M+H]^+$ 1406.8411 | $[M+H]^+$ 1406.8480 | 4.90 |
| **BNP37C2-L8** | C69H111N15O16 | $[M+H]^+$ 1406.8411 | $[M+H]^+$ 1406.8474 | 4.48 |
| **BNP37C2-L9** | C68H97N15O17 | $[M+H]^+$ 1396.7265 | $[M+H]^+$ 1396.7308 | 3.07 |
| **BNP37C2-L10** | C68H98N16O16 | $[M+H]^+$ 1395.7425 | $[M+H]^+$ 1395.7487 | 4.44 |
| **BNP37C2-L11** | C68H97N17O16 | $[M+H]^+$ 1408.7377 | $[M+H]^+$ 1408.7443 | 4.69 |
| **BNP37C2-L12** | C72H101N15O16 | $[M+H]^+$ 1432.7629 | $[M+H]^+$ 1432.7694 | 4.53 |
| **BNP37C2-L13** | C74H101N15O16 | $[M+H]^+$ 1456.7629 | $[M+H]^+$ 1456.7712 | 5.70 |
| **BNP37C2-L14** | C65H97N17O16 | $[M+H]^+$ 1372.7377 | $[M+H]^+$ 1372.7445 | 4.95 |
| **BNP37C2-L15** | C73H99N15O17 | $[M+H]^+$ 1458.7422 | $[M+H]^+$ 1458.7489 | 4.59 |
| **BNP37C2-L16** | C73H99N15O17 | $[M+2H]^{2+}$ 729.8750 | $[M+2H]^{2+}$ 729.8748 | -0.27 |
| **BNP37C2-L17** | C72H101N15O16 | $[M+H]^+$ 1432.7629 | $[M+H]^+$ 1432.7706 | 5.37 |
| **BNP37C2-L18** | C71H106N16O16 | $[M+2H]^{2+}$ 720.4065 | $[M+2H]^{2+}$ 720.4065 | 0.00 |
| **BNP37C2-L19** | C70H104N16O17 | $[M+2H]^{2+}$ 721.3961 | $[M+2H]^{2+}$ 721.3959 | -0.27 |
| **BNP37C2-L20** | C71H107N17O16 | $[M+2H]^{2+}$ 727.9119 | $[M+2H]^{2+}$ 727.9122 | -0.41 |
| **BNP37C2-L21** | C71H107N17O16 | $[M+2H]^{2+}$ 727.9119 | $[M+2H]^{2+}$ 727.9131 | 1.65 |
| **BNP37C2-L22** | C66H97N15O16 | $[M+2H]^{2+}$ 678.8679 | $[M+2H]^{2+}$ 678.8687 | -1.46 |
| **BNP37C2-L23** | C83H115N15O17 | $[M+H]^+$ 1594.8674 | $[M+H]^+$ 1595.8799 | 7.84 |
| **BNP37C2-L24** | C70H112N15016 | $[M+H]^+$ 1420.8568 | $[M+H]^+$ 1420.8622 | 3.80 |
| **BNP37C2-L25** | C72H117N15O16 | $[M+H]^+$ 1448.8881 | $[M+H]^+$ 1448.8978 | 6.69 |
| **BNP37C2-L26** | C74H121N15O16 | $[M+H]^+$ 1476.9194 | $[M+H]^+$ 1476.9280 | 5.82 |
| **BNP37C2-L27** | C76H125N15O16 | $[M+H]^+$ 1504.9507 | $[M+H]^+$ 1504.9622 | 7.64 |
| **BNP37C2-L28** | C73H111N15O16 | $[M+H]^+$ 1454.8411 | $[M+H]^+$ 1454.8464 | 3.64 |
| **BNP37C2-L29** | C67H96N16O16 | $[M+H]^+$ 1381.7268 | $[M+H]^+$ 1381.7327 | 4.27 |
| **BNP37C2-L30** | C72H100FN15O16 | $[M+H]^+$ 1450.7535 | $[M+H]^+$ 1450.7582 | 3.23 |
| **BNP37C2-L31** | C72H107N15O16 | $[M+H]^+$ 1438.8098 | $[M+H]^+$ 1438.8124 | 1.81 |
| **BNP37C2-L32** | C69H101N15O16 | $[M+H]^+$ 1396.7629 | $[M+H]^+$ 1396.7668 | 2.79 |
| **BNP37C2-L33** | C66H95Cl2N15O16 | $[M+H]^+$ 1424.6537 | $[M+H]^+$ 1424.6593 | 3.93 |

| | | | | |
|---|---|---|---|---|
| **BNP37C2-L34** | C67H99N15O16 | $[M+H]^+$ 1370.7472 | $[M+H]^+$ 1370.7566 | 6.85 |
| **BNP37C2-L35** | C66H96ClN15O16 | $[M+H]^+$ 1390.6926 | $[M+H]^+$ 1390.7010 | 6.04 |
| **BNP37C2-L36** | C66H96BrN15O16 | $[M+H]^+$ 1434.6421 | $[M+H]^+$ 1434.6521 | 6.97 |
| **BNP37C2-L37** | C66H96FN15O16 | $[M+H]^+$ 1374.7222 | $[M+H]^+$ 1374.7306 | 6.11 |
| **BNP37C2-Ala1** | C72H116N14O16 | $[M+H]^+$ 1433.8772 | $[M+H]^+$ 1433.8741 | -2.16 |
| **BNP37C2-Ala3** | C72H116N14O16 | $[M+H]^+$ 1433.8772 | $[M+H]^+$ 1433.8750 | -1.53 |
| **BNP37C2-Ala4** | C70H113N15O16 | $[M+H]^+$ 1420.8568 | $[M+H]^+$ 1420.8549 | -1.53 |
| **BNP37C2-Ala5** | C72H116N14O16 | $[M+H]^+$ 1433.8772 | $[M+H]^+$ 1433.8755 | -1.18 |
| **BNP37C2-Ala6** | C67H115N15O16 | $[M+H]^+$ 1386.8724 | $[M+H]^+$ 1386.8700 | -1.73 |
| **BNP37C2-Ala7** | C70H113N15O16 | $[M+H]^+$ 1420.8568 | $[M+H]^+$ 1420.8553 | -1.05 |
| **BNP37C2-Ala8** | C72H116N14O16 | $[M+H]^+$ 1433.8772 | $[M+H]^+$ 1433.8751 | -1.46 |
| **BNP37C2-Ala9** | C67H115N15O15 | $[M+H]^+$ 1370.8775 | $[M+H]^+$ 1370.8755 | -1.45 |
| **BNP37C2-Ala10** | C71H115N15O16 | $[M+H]^+$ 1434.8724 | $[M+H]^+$ 1434.8690 | -2.37 |
| **BNP37C2-Ala11** | C72H119N15O14 | $[M+H]^+$ 1418.9139 | $[M+H]^+$ 1438.9121 | -1.27 |
| **BNP37C2-Dab4** | C71H116N16O16 | $[M+H]^+$ 1449.8833 | $[M+H]^+$ 1449.8889 | 3.86 |
| **BNP37C2-Dab6** | C68H118N16O16 | $[M+H]^+$ 1415.8990 | $[M+H]^+$ 1415.9056 | 4.66 |
| **BNP37C2-Dab7** | C71H116N16O16 | $[M+2H]^{2+}$ 725.4456 | $[M+2H]^{2+}$ 725.4484 | 3.86 |
| **BNP37C2-Dab9** | C68H118N16O15 | $[M+H]^+$ 1399.9041 | $[M+H]^+$ 1399.9075 | 1.86 |
| **BNP37C2-Dab10** | C72H118N16O16 | $[M+H]^+$ 1463.8990 | $[M+H]^+$ 1463.9054 | 4.37 |
| **BNP37C2-Dab11** | C73H122N16O14 | $[M+H]^+$ 1447.9405 | $[M+H]^+$ 1447.9447 | 2.90 |

| | Pathogens | Strain Description | Resistant phenotype | MIC (μg/mL) | | | | | | | | | | |
|---|---|---|---|---|---|---|---|---|---|---|---|---|---|---|
| | | | | Amoxicillin | Ampicillin | Azithromycin | Aztreonam | Ceftizoxime | Ciprofloxacin | Colistin | Gentamicin | Imipenam | Meropenem | Tetracycline |
| **Gram-** | *K. pneumoniae* | ATCC13883 | Amx$^R$, Amp$^R$,Cip$^R$ | >64 | >64 | 1 | 0.0625 | 0.125 | 16 | 0.25 | 2 | 4 | 0.125 | 1 |
| | *K. pneumoniae* | BNCC359393 | Amx$^R$, Amp$^R$, Azm$^R$, Cza$^R$, Cip$^R$, Col$^R$, Gen$^R$, Ipm$^R$, Mer$^R$ | >64 | >64 | >64 | 0.0625 | >64 | >64 | >64 | >64 | >64 | 64 | 4 |
| | *A. baumannii* | BNCC194496 | Cip$^R$ | 8 | 16 | 1 | 16 | 4 | 16 | 0.25 | 4 | 0.5 | 0.25 | 2 |
| | *A. baumannii* | ATCC19606 | Amx$^R$, Amp$^R$, Cip$^R$ | >64 | >64 | 1 | 16 | 16 | 32 | 0.25 | 16 | 1 | 0.5 | 4 |
| | *A. baumannii* | ATCCBAA-1605 | Amx$^R$, Amp$^R$, Cza$^R$, Cip$^R$, Gen$^R$, Mer$^R$, Ipm$^R$, Tcy$^R$ | >64 | >64 | 16 | >64 | >64 | >64 | 0.25 | >64 | >64 | 16 | >64 |
| | *P. aeruginosa* | ATCC27853 | Amx$^R$, Amp$^R$, Cip$^R$ | >64 | 64 | 32 | 2 | 16 | 32 | 0.5 | 2 | 4 | 0.25 | 16 |
| | *E. cloacae* | ATCC13047 | Amx$^R$, Col$^R$,Cza$^R$ | >64 | 64 | 4 | 0.5 | 2 | 1 | >64 | 4 | 2 | 0.125 | 2 |
| | *E. cloacae* | BNCC185928 | Azm$^R$, Col$^R$,Cza$^R$ | 4 | 64 | 32 | 2 | 4 | 2 | >64 | 4 | 2 | 0.125 | 2 |
| **Gram+** | | | | Amoxillin | Azithromycin | Erythromycin | Lincomycin | Linozolid | Methicillin | Penicillin | Vancomycn | | | |
| | *S. aureus* | BAA44 | Amx$^R$, Azm$^R$, Ery$^R$ , Lin$^R$, Met$^R$, Pen$^R$ | 64 | >64 | >64 | >64 | 0.5 | >64 | 32 | 1 | | | |
| | *S. aureus* | BNCC 186335 | Azm$^R$, Pen$^R$ | 0.5 | 8 | 4 | 4 | 0.5 | 0.25 | 2 | 0.5 | | | |
| | *S. aureus* | USA300 | Amx$^R$, Azm$^R$, Ery$^R$, Met$^R$, Pen$^R$ | 64 | >64 | >64 | 2 | 1 | >64 | 32 | 1 | | | |
| | *S. pyogenes* | 8007 | Ery$^R$ | - | - | >64 | - | - | - | - | 0.5 | | | |
| **Fungi** | | | | Caspofungin | Fluconazole | Terbinafine | 5-Fluorocytosine | | | | | | | |
| | *C. alicans* | BNCC 186382 | Fca$^R$ | ＜0.125 | 4 | 4 | 8 | | | | | | | |
| | *A. fumigatus* | BNCC 122691 | Fca$^R$, 5-FU$^R$, Tbf$^R$ | 2 | >64 | 16 | 32 | | | | | | | |

Note: Amp$^R$: Ampicillin resistance; Amx$^R$: Amoxicillin resistance; Azm$^R$: Azithromycin resistance; Cip$^R$: Ciprofloxacin resistance Col$^R$: Colistin resistance; Cza$^R$: Ceftizoxime resistace; Ery$^R$: Erythromycin resistance; FCA$^R$: Fluconazole resistance; Gen$^R$: Gentamycin resistance; Ipm$^R$: Imipenem resistance; Mer$^R$: Meropenem resistance; Lin$^R$: Lincomycin resistance; Met$^R$ : Methicillin resistance; Pen$^R$: Penicillin resistance; Tbf$^R$: Terbinafine resistance; Tcy$^R$: Tetracycline resistance; 5-FU$^R$: 5-Fluorocytosine resistance.

**Table S4:** MIC values of drug-resistant strains used in this study.

**Table S5:** Pharmacokinetic parameters of paenimycin and colistin (n= 3 rats per group). One rat died in the colistin group via i.v. injection.

| Injection method | PK parameters | Paenimycin (Mean ± SD) | Colisin (Mean ± SD) |
|---|---|---|---|
| s.c. (10 mg/kg) | $T_{1/2}$ (h) | 20.2 ± 1.9 | 2.04 ± 0.03 |
| | $T_{max}$ (h) | 8 ± 0 | 2.67 ± 1.15 |
| | $C_{max}$ (ng/mL) | 5200 ± 832 | 2753 ± 285 |
| | $AUC_{last}$ (h*ng/mL) | 113551 ± 14845 | 22458 ± 4769 |
| | $AUC_{Inf}$ (h*ng/mL) | 115251 ± 14868 | 22458 ± 4769 |
| | $AUC_{_\%Extrap_}$obs (%) | 1.49 ± 0.18 | 0.0486 ± 0.0019 |
| | $MRT_{Inf_}$obs (h) | 18.0 ± 0.6 | 4.88 ± 0.31 |
| | $AUC_{last}$/D (h*mg/mL) | 11355 ± 1484 | 2246 ± 477 |
| | F (%) | 102 ± 13 | 90 ± 19 |
| i.v. (5 mg/kg) | Cl_obs (mL/min/kg) | 1.48 ± 0.17 | 6.81 |
| | $T_{1/2}$ (h) | 8.28 ± 0.94 | 2.85 |
| | $C_0$ (ng/mL) | 28111 ± 4675 | 15356 |
| | $AUC_{last}$ (h*ng/mL) | 56303 ± 6510 | 12497 |
| | $AUC_{Inf}$ (h*ng/mL) | 56747 ± 6337 | 12506 |
| | $AUC_{_\%Extrap_}$obs (%) | 0.81 ± 0.42 | 0.0795 |
| | $MRT_{Inf_}$obs (h) | 4.82 ± 0.58 | 1.83 |
| | $AUC_{last}$/D (h*mg/mL) | 11261 ± 1302 | 2499 |
| | $V_{ss_}$obs (L/kg) | 0.432 ± 0.101 | 0.736 |

**Table S6:** Bacterial strains, human cells lines and culture conditions.

| | **Organism** | **Strain name** | **Media** | **Culture condition** |
|---|---|---|---|---|
| **Gram+** | *Staphylococcus aureus* | BNCC 186335 | LB | 37 ℃, aerobic |
| | *Staphylococcus aureus* | ATCC BAA44 | LB | 37 ℃, aerobic |
| | *Streptococcus pyogenes* | BNCC 336670 | BHI-MM | 37 ℃, 5%$CO_2$ |
| | *Streptococcus pyogenes* | 8007 | BHI-MM | 37 ℃, 5%$CO_2$ |
| **Gram-** | *Klebsiella pneumoniae* | BNCC 359393 | LB | 37 ℃, aerobic |
| | *Klebsiella pneumoniae* | ATCC 13883 | LB | 37 ℃, aerobic |
| | *Acinetobacter baumannii* | ATCC 19606 | LB | 37 ℃, aerobic |
| | *Acinetobacter baumannii* | ATCC BAA1605 | LB | 37 ℃, aerobic |
| | *Acinetobacter baumannii* | BNCC 194496 | LB | 37 ℃, aerobic |
| | *Acinetobacter baumannii* | BNCC 194496-*mcr-1* | LB | 37 ℃, aerobic |
| | *Pseudomonas aeruginosa* | ATCC 27853 | LB | 37 ℃, aerobic |
| | *Enterobacter cloacae* | BNCC 185928 | LB | 37 ℃, aerobic |
| | *Enterobacter cloacae* | ATCC 13047 | LB | 37 ℃, aerobic |
| | *Escherichia coli* | ATCC25922 | LB | 37 ℃, aerobic |
| | *Escherichia coli* | clinical isolate 9 | LB | 37 ℃, aerobic |
| | *Escherichia coli* | clinical isolate 18 | LB | 37 ℃, aerobic |
| | *Escherichia coli* | MG1655 | LB | 37 ℃, aerobic |
| | *Escherichia coli* | MG1655-*mcr-1* | LB | 37 ℃, aerobic |
| | *Neisseria gonorrhoeae* | BNCC 270346 | GC | 37 ℃, 5%$CO_2$ |
| | *Neisseria meningitidis* | BNCC 353914 | GC | 37 ℃, 5%$CO_2$ |
| Fungi | *Candida albicans* | BNCC 186382 | YPD | 30 ℃, aerobic |
| | *Aspergillus fumigatus* | BNCC 122691 | YPD | 30 ℃, aerobic |
| Human cells | HepG2 | TCHu72 | DMEM+10%FBS | 37 ℃, 5%$CO_2$ |
| | HK-2 | ATCC-CRL-1469 | RPMI-1640+10%FBS | 37 ℃, 5%$CO_2$ |
